# Independent regulation of early trafficking of NMDA receptors by ligand-binding domains of the GluN1 and GluN2A subunits

**DOI:** 10.1101/2024.02.02.578575

**Authors:** Jakub Netolicky, Petra Zahumenska, Anna Misiachna, Marharyta Kolcheva, Tomas Kucera, Jakub Fibigar, Kristyna Rehakova, Katarina Hemelikova, Stepan Kortus, Emily Langore, Marek Ladislav, Jan Korabecny, Martin Horak

## Abstract

The essential role of N-methyl-D-aspartate receptors (NMDARs) in excitatory neurotransmission is underscored by numerous pathogenic variants in the GluN subunits, including those identified in their ligand-binding domains (LBDs). The prevailing hypothesis postulates that the endoplasmic reticulum (ER) quality control machinery verifies the agonist occupancy of NMDARs; however, whether it controls the structure of LBDs or the functionality of NMDARs is unknown. Using alanine substitutions combined with microscopy and electrophysiology, we found that surface expression of GluN1/GluN2A receptors, the primary NMDAR subtype in the adult forebrain, strongly correlates with EC_50_ values for glycine and L-glutamate. Interestingly, co-expression of both GluN1 and GluN2A subunits with alanine substitutions led to an additive reduction in the surface number of GluN1/GluN2A receptors, as did co-expression of both GluN1 and GluN2A subunits containing closed cleft conformation of LBDs. The synchronized ER release confirmed the altered regulation of early trafficking of GluN1/GluN2A receptors bearing alanine substitutions in the LBDs. Furthermore, the human versions of GluN1/GluN2A receptors containing pathogenic GluN1-S688Y, GluN1-S688P, GluN1-D732E, GluN2A-S511L, and GluN2A-T690M variants exhibited distinct surface expression compared to the corresponding alanine substitutions. Mutant cycles of GluN1-S688, GluN1-D732, GluN2A-S511, and GluN2A-T690 residues revealed, in most cases, a weak correlation between surface expression of the mutant GluN1/GluN2A receptors and their EC_50_ values for glycine or L-glutamate. Consistent with our experimental data, molecular modeling and dynamics showed that the ER quality control machinery likely perceives structural changes of the LBDs but not the functionality of GluN1/GluN2A receptors.

**Significant statement:** Our study showed that structural changes in LBDs independently regulate the early trafficking of GluN1/GluN2A receptors and that the surface numbers of mutant GluN1/GluN2A receptors do not necessarily correlate with their agonist sensitivity. In addition, we validated a novel system of synchronized release of GluN1/GluN2A receptors from the ER. Together, our experimental and *in silico* findings support the urgency of further detailed research on the regulation of early trafficking of NMDARs, as it may open the avenue to targeted therapeutic intervention of CNS disorders associated with pathogenic variants in GluN subunits.

## Introduction

N-methyl-D-aspartate receptors (NMDARs) are a family of ionotropic glutamate receptors that play a crucial role in excitatory transmission in the mammalian central nervous system (CNS). Conventional NMDARs are heterotetramers of two GluN1 and two GluN2 subunits in a 1-2-1-2 arrangement (Karakas and Furukawa, 2014). Eight known splice variants of the GluN1 subunit arise from a single gene, and there are four different genes for the GluN2 subunits: GluN2A, GluN2B, GluN2C, and GluN2D (Sanz-Clemente et al., 2013; Vieira et al., 2020; Hansen et al., 2021). Each GluN subunit consists of four membrane helices (M1-M4), an intracellular C-terminal domain (CTD), an extracellular amino-terminal domain (ATD), and an extracellular loop between M3 and M4 (Figure 1A). The ligand-binding domain (LBD) is formed by two segments, S1 and S2 (Traynelis et al., 2010; Paoletti et al., 2013). Critical amino acid residues in the LBDs that are involved either by direct interactions or via water molecules in the interaction with agonists (glycine in the GluN1 subunit and L-glutamate in the GluN2 subunits, respectively) are highly conserved among mammals, highlighting their critical importance for the proper function of NMDARs (Stroebel and Paoletti, 2021).

**Figure 1.**
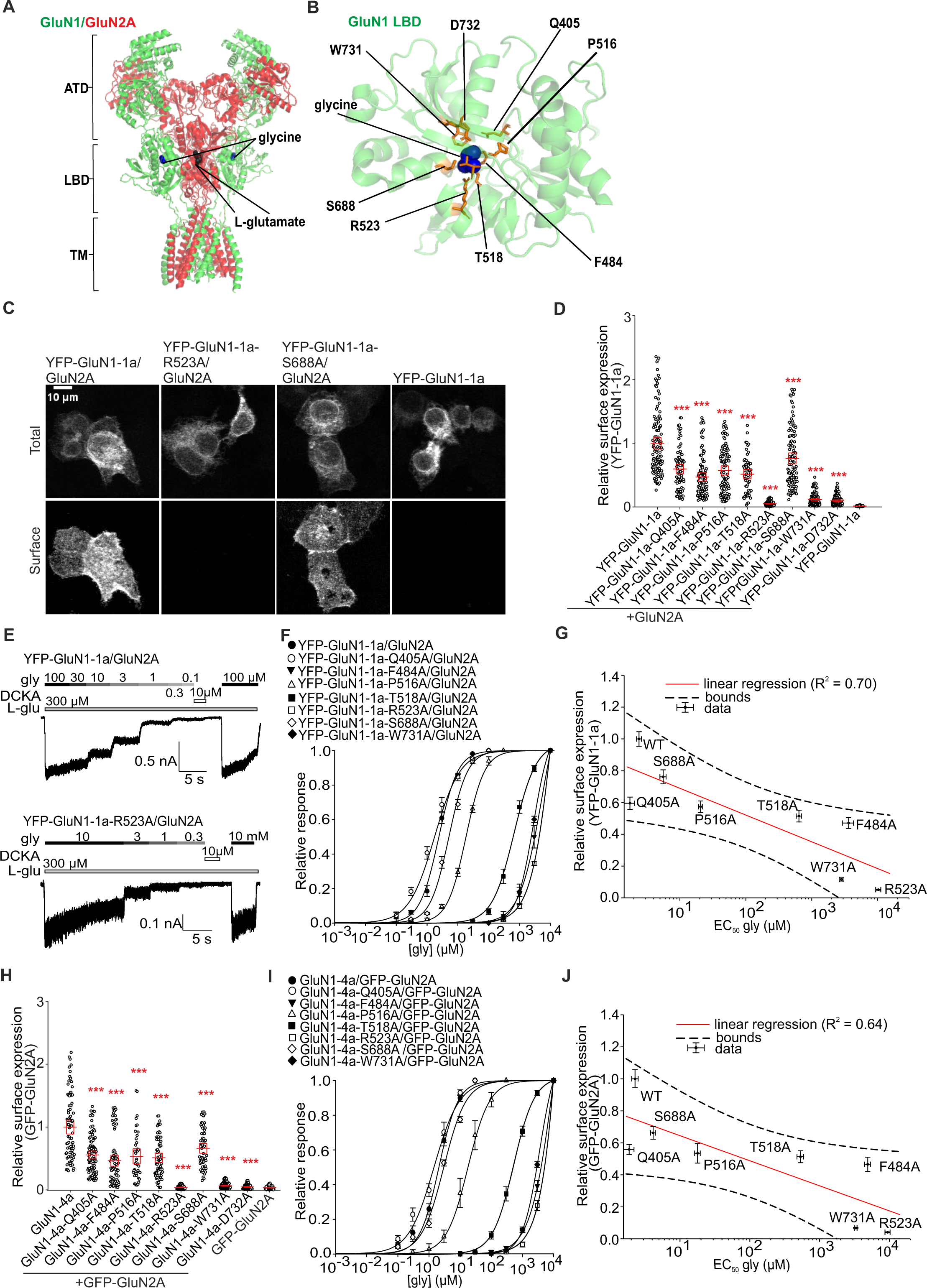
Mutations of residues in the LBD of the GluN1 subunit that interact with glycine affect surface expression and glycine potency of GluN1/GluN2A receptors. **(A)** The structural model of the NMDAR comprises GluN1 (depicted in green) and GluN2A (depicted in red) subunits (based on PDB ID: 7EU7). (**B**) Structural model of the LBD of GluN1 subunit with labeled glycine-interacting amino acids (based on PDB ID: 1PB7). (**C**) Representative images of HEK293 cells transfected with either the YFP-GluN1-1a subunit (WT or mutant variant) alone (negative control) or with the GluN2A subunit. The total and surface signals (top and bottom row, respectively) of YFP-GluN1-1a subunits were labeled using an anti-GFP antibody 24 hours after the transfection. (**D,H**) Summary of the relative surface expression of NMDARs consisting of either WT or mutated YFP-GluN1-1a subunit co-expressed with the GluN2A subunit (**D**) and WT or mutated GluN1-4a subunit co-expressed with the GFP-GluN2A subunit (**H**), measured using fluorescence microscopy; \*\*\**p* < 0.001 vs WT; one-way ANOVA. Data points correspond to individual cells (*n* ≥ 56), and the red box plot represents mean ± SEM. (**E**) Representative whole-cell patch-clamp recordings of HEK293 cells expressing the indicated NMDAR subunits. Glycine (gly) at the indicated concentrations was applied in the continuous presence of 300 µM L-glutamate (L-glu). (**F, I**) Normalized concentration-response curves for glycine measured from HEK293 cells expressing NMDARs containing YFP-GluN1-1a (**F**) or GluN1-4a subunit variants (**I**) co-expressed with the GluN2A subunit were obtained by fitting the data using Equation (1) (see Methods); for the summary of fitting parameters see **Table 1**. (**G,J**) Correlation of surface expression and glycine EC_50_ values at NMDARs composed of WT or mutated YFP-GluN1-1a subunit (**G**) and WT or mutated GluN1-4a (**J**) subunit expressed together with GFP-GluN2A subunit; data were fitted by linear regression.

**Table 1.** Summary of fitting parameters for steady-state concentration-response curves for glycine effect measured in HEK293 cells expressing the indicated NMDAR subunits carrying a mutation in the LBD of the GluN1 subunit.

| Receptor | $EC_{50}$ (μM) | $h$ | $n$ | Receptor | $EC_{50}$ (μM) | $h$ | $n$ |
| --- | --- | --- | --- | --- | --- | --- | --- |
| YFP-GluN1-1a/<br>GluN2A | $2.40 \pm 0.23$ | $1.41 \pm 0.03$ | 5 | GluN1-4a/<br>GFP-GluN2A | $2.27 \pm 0.26$ | $1.31 \pm 0.15$ | 8 |
| YFP-GluN1-1a-Q405A/<br>GluN2A | $1.75 \pm 0.34$ | $1.04 \pm 0.08$ | 8 | GluN1-4a-Q405A/<br>GFP-GluN2A | $1.87 \pm 0.36$ | $0.95 \pm 0.09$ | 5 |
| YFP-GluN1-1a-F484A/<br>GluN2A | $3,624.55 \pm 787.45$ | $1.50 \pm 0.13$ | 4 | GluN1-4a-F484A/<br>GFP-GluN2A | $4,897.02 \pm 512.26$ | $1.62 \pm 0.14$ | 6 |
| YFP-GluN1-1a-P516A/<br>GluN2A | $20.61 \pm 1.38$ | $1.29 \pm 0.04$ | 7 | GluN1-4a-P516A/<br>GFP-GluN2A | $17.87 \pm 1.77$ | $1.25 \pm 0.05$ | 9 |
| YFP-GluN1-1a-T518A/<br>GluN2A | $640.72 \pm 63.64$ | $1.44 \pm 0.03$ | 7 | GluN1-4a-T518A/<br>GFP-GluN2A | $533.11 \pm 62.20$ | $1.36 \pm 0.06$ | 8 |
| YFP-GluN1-1a-R523A/<br>GluN2A | $10,261.34 \pm 1,052.06$ | $1.31 \pm 0.08$ | 4 | GluN1-4a-R523A/<br>GFP-GluN2A | $9,451.12 \pm 886.52$ | $1.61 \pm 0.10$ | 5 |
| **YFP-GluN1-1a-S688A/**<br>**GluN2A** | 5.48 ± 0.56 | 1.32 ± 0.06 | 5 | **GluN1-4a-S688A/**<br>**GFP-GluN2A** | 4.12 ± 0.35 | 1.03 ± 0.08 | 7 |
| **YFP-GluN1-1a-W731A/**<br>**GluN2A** | 2,824.15 ± 183.58 | 1.61 ± 0.07 | 5 | **GluN1-4a-W731A/**<br>**GFP-GluN2A** | 3,296.95 ± 165.39 | 1.72 ± 0.02 | 6 |
| **YFP-GluN1-1a-D732A/**<br>**GluN2A** | NR | - | 12 | **GluN1-4a-D732A/**<br>**GFP-GluN2A** | NA | - | 10 |

The EC<sub>50</sub> and *h* values were obtained by fitting the data from individual cells to Equation (1). There is no significant difference between YFP-GluN1-1a and GluN1-4a versions; *p*>0.050, Student's *t*-test. NR indicates “not responding”, and NA indicates “not analyzed”.

One of the critical parameters contributing to the physiological or pathophysiological roles of NMDARs is their number on the cell surface, determined by the balance between their exocytosis and endocytosis. Exocytosis (hereafter referred to as “early trafficking”) of NMDARs is regulated at several levels. This includes control over their translation, the proper assembly of functional heterotetramers and maturation in the endoplasmic reticulum (ER), and intracellular transport via Golgi apparatus (GA) (Meddows et al., 2001; Schüler et al., 2008; Horak et al., 2014; Hanus and Ehlers, 2016). Alternative pathways bypassing the Golgi apparatus have also been identified (Jeyifous et al., 2009). The prevailing hypothesis dealing with the processing of NMDARs in the ER postulates that the ER quality control machinery verifies the sensitivity of NMDARs to their (co)-agonists via a common mechanism shared with the α-amino-3-hydroxy-5-methyl-4-isoxazolepropionic acid receptors (AMPARs) and kainate receptors (KARs) (Penn et al., 2008; Coleman et al., 2009, 2010; Scholefield et al., 2019; Horak et al., 2021). However, in the case of GluN1/GluN2 receptors, this hypothesis is supported by a relatively limited number of studies, namely (i) on the mutant GluN1-D732A/GluN2A receptor, which shows virtually zero surface expression and an extremely high EC_50_ value for glycine due to disruption of direct glycin interaction with GluN1-D732 residue (Kenny et al., 2009), (ii) GluN1/GluN2B receptors carrying mutations that disrupt direct (GluN2B-E413A, GluN2B-S690G, GluN2B-R519K) and water (GluN2B-V686A) interaction with L-glutamate or that indirectly lead to changes in EC_50_ values for L-glutamate (GluN2B-F416S) (She et al., 2012), and (iii) pathogenic variants in the LBDs of the GluN2A and GluN2B subunits, specifically, that disrupt direct (GluN2B-E413G, GluN2A-R518H) and water (GluN2A-G483R, GluN2A-V685G, GluN2A-D731N) interaction with L-glutamate or are present elsewhere in the LBDs (GluN2A-T531M, GluN2B-C461F, GluN2A-I694T, GluN2A-M705V, GluN2A-A716T, GluN2A-A727T, GluN2A-V734L, GluN2A-K772E) (Swanger et al., 2016). However, our recent study showed that even a profound reduction in the EC_50_ value for glycine does not lead to a reduction in the surface number of GluN1-S688Y/GluN2 receptors (Skrenkova et al., 2020), supporting further research addressing the roles of LBDs in the early trafficking of GluN1/GluN2 receptors.

We investigated the early trafficking of GluN1/GluN2A receptors by mutagenesis of amino acid residues in the LBDs of GluN1 and GluN2A subunits involved in direct interaction with agonists. Our experiments showed a strong (for alanine substitutions in the GluN1 subunit) or weak (for alanine substitution in the GluN2A subunit) correlation between surface expression levels and EC_50_ values for agonists. In addition, co-expression of both GluN1 and GluN2A subunits with alanine substitutions additively decreased the surface numbers of GluN1/GluN2A receptors. The findings from the study of pathogenic variants, mutant cycles, and molecular dynamics essentially refute the assumption of a correlation between trafficking and agonist potency and point to a complex role of structural changes in regulating early trafficking of GluN1/GluN2A receptors.

## Materials and Methods

### Molecular biology

We used the following DNA vectors carrying genes encoding rat versions of the YFP-GluN1-1a (NP_058706.1), GluN1-4a (NP_001257539.1), GluN2A or GFP-GluN2A (NP_036705), GluN2B or GFP-GluN2B (NP_036706), GluN3A (NP_612555.1), human GluN2A (hGluN2A, NP_001127879.1) subunits. Human versions of YFP-GluN1-1a (YFP-hGluN1-1a) and GluN1-4a (hGluN1-4a) were prepared as we published previously (Skrenkova et al., 2020). The closed cleft conformation of LBDs of the GluN1-4a (GluN1-4a-N499C-Q686C) and GFP-GluN2A (GFP-GluN2A-K487C-N687C) subunits were constructed according to previous studies (Blanke and VanDongen, 2008; Kussius and Popescu, 2010; Dai and Zhou, 2016). An ARIAD-mNEONGreen-GluN1-1a construct (ARIAD-mNEON-GluN1-1a) containing the ARIAD sequence (Hangen et al., 2018) followed by a rat version of the GluN1-1a subunit with an inserted mNEONGreen sequence after the 21st amino acid residue was prepared by gene synthesis (Thermo Fischer Scientific) and subsequently subcloned into the pcDNA3 expression vector. Point amino acid substitutions in genes encoding GluN subunits were performed using the QuikChange site-directed mutagenesis kit (Agilent Technologies) and verified by DNA sequencing.

### HEK293 cells and primary hippocampal neurons

Human embryonic kidney 293 (HEK293) cells were grown in Opti-MEM I medium supplemented with 5% fetal bovine serum (FBS; both from Thermo Fischer Scientific) and used in experiments up to passage 10. For electrophysiology, HEK293 cells in 24-well plates were transfected with a total of 0.9 μg of DNA vectors carrying genes encoding GluN1 and GluN2 subunits and green fluorescent protein (GFP; pQBI 25 vector; for identification of transfected cells; Takara) in a 1:1:1 ratio using 0.9 μl of PolyMag reagent (OZBiosciences) in 50 μl of Opti-MEM I medium. After 20 min incubation on a magnetic plate, HEK293 cells were trypsinized and resuspended in Opti-MEM I medium containing 1% FBS, 2 mM MgCl_2_, and 10 µM 5,7-dichlorokynurenic acid (DCKA, HelloBio; in the case of NMDARs with mutant GluN2 subunit) or 10 µM D-2-amino-5-phosphonopentanoic acid (D-APV, HelloBio; in the case of NMDARs with mutant GluN1 subunit), to reduce cell death caused by excitotoxicity. For microscopy, HEK293 cells were grown in 12-well plates and transfected with a total of 0.45 μg of DNA vectors carrying genes encoding GluN1 and GluN2 subunits at a ratio of 1:2 (YFP-/GFP-GluN subunit versus untagged GluN subunit), using 1 μl of Lipofectamine 2000 reagent (Thermo Fischer Scientific) in 50 μl of Opti-MEM I medium. DNA vectors carrying wild-type (WT) or mutant ARIAD-mNEON-GluN1-1a constructs were co-transfected at a 1:2 ratio with untagged GluN2A or GluN3A subunits.

All animal procedures were performed in accordance with the ARRIVE guidelines and the European Commission Council Directive 2010/63/EU for animal experiments. Primary cultures of hippocampal neurons were prepared from Wistar rats of both sexes (at embryonic day 18) by dissecting hippocampal neurons in a cold dissection solution consisting of Hanks’ balanced salt solution supplemented with 10 mM HEPES (pH 7.4). Hippocampi were then incubated in a dissection medium containing 0.05% trypsin and 0.1 mg/ml DNase I (Merck) for 20 min at 37°C. Dissociated cells were plated on poly-L-lysine-coated glass coverslips at a density of 2 × 10,000 cells per cm^2^ in a plating medium consisting of minimal essential medium (MEM) supplemented with 10% heat-inactivated horse serum, N2 supplement (1x), 1 mM sodium pyruvate, 20 mM D-glucose, 25 mM HEPES, and 1% penicillin-streptomycin. After 3 hours, the plating medium was replaced entirely with Neurobasal medium enriched with 2% B-27 and 2 mM L-glutamine (all purchased from Thermo Fischer Scientific). Every 3-4 days, ∼50% of the volume of this culture medium was replaced with fresh medium. Hippocampal neurons were transfected at day 6 *in vitro* (DIV6) with DNA vectors carrying genes encoding the indicated GluN subunits using Lipofectamine 2000 (Lichnerova et al., 2015)

### Immunofluorescence microscopy

Immunofluorescence labeling was performed 24 hours after the completion of transfection of HEK293 cells. Hippocampal neurons were labeled 48 hours after the completion of transfection (DIV14). All primary and secondary antibodies were diluted in a blocking solution consisting of PBS and 0.2% (v/v) bovine serum albumin (BSA, Merck). Cells were first washed with ice-cold PBS and incubated for 5 min in a blocking solution on ice. Surface antigens were labeled with primary antibody (rabbit anti-GFP, 1:1000, Merck) for 15 min on ice, and then cells were washed with ice-cold blocking solution and incubated with secondary antibody (goat anti-rabbit with Alexa Fluor 555, 1:1000, Thermo Fisher Scientific). Cells were then washed with ice-cold PBS, fixed in 4% paraformaldehyde (PFA) in PBS for 15 min, permeabilized with 0.25% Triton X-100 in PBS for 5 min, and incubated with primary antibody (mouse anti-GFP, 1:1000, Neuromab), and then with secondary antibody (goat anti-mouse conjugated with Alexa Fluor 488, 1:1000, Thermo Fisher Scientific) to label intracellular antigens. In experiments with ARIAD-mNEON-GluN1-1a constructs, HEK293 cells were first incubated with ARIAD ligand at a concentration of 1 µM (AL; D/D Solubilizer; Takara) for the indicated times, fixed with PFA for 7 min, labeled using primary antibody (mouse anti-mNEONGreen, 1:1000, ChromoTek) and secondary antibody (goat anti-mouse conjugated with Alexa Fluor 555, 1:1000, Thermo Fisher Scientific) and re-fixed by PFA; intracellular mNEONGreen epitopes in HEK293 cells were not labeled with antibodies. To detect the cis-GA structures, we used fixed HEK293 cells labeled with primary antibody (rabbit anti-GM130, 1:1000, Merck) and secondary antibody (goat anti-rabbit conjugated with Alexa Fluor 555, 1:1000; labeled cells were then re-fixed with PFA in PBS for 5 min. Microscopic samples were mounted on glass slides using ProLong Antifade reagent (Thermo Fisher Scientific). Images were captured using an Olympus FV10i microscope with a 60x/1.35 oil immersion objective and analyzed using ImageJ 1.52N software (National Institutes of Health). In the case of HEK293 cells, total and surface fluorescence intensity was analyzed on whole cells; in the case of hippocampal neurons, 10 separate 5-µm segments of secondary or tertiary dendrites from a single neuron were analyzed. The degree of co-localization of GM130 and mNEONGreen signals was determined as the ratio between the intensity of the mNEONGreen signal co-localizing with the GM130 signal and the intensity of the mNEONGreen signal outside the co-localization region as used previously (Skrenkova et al., 2019). The following competitive antagonists were employed for microscopical experiments: trans-2-Carboxy-5,7-dichloro-4-phenylaminocarbonylamino-1,2,3,4-tetrahydroquinoline (L-689-560; Tocris); 7-Chloro-4-hydroxy-3-(3-phenoxy)phenyl-2(1H)-quinolinone (L-701-324; Tocris); (R)-3-(2-Carboxypiperazin-4-yl)propyl-1-phosphonic acid ((R)-CPP; Merck); [[[(1S)-1-(4-Bromophenyl)ethyl]amino](1,2,3,4-tetrahydro-2,3-dioxo-5-quinoxalinyl)methyl] phosphonic acid tetrasodium salt (PEAQX; HelloBio).

### Electrophysiology

Whole-cell current responses of HEK293 cells were recorded by the patch-clamp method ∼24-48 hours after completion of transfection using an Axopatch 200B amplifier (Axon Instruments) with series resistance (<10 MΩ) and capacitance compensation (∼80%). Current responses of NMDARs were recorded in voltage-clamp mode with constant membrane potential held at -60 mV, filtered using an eight-pole Bessel filter (frequency < 2 kHz), sampled at 5 kHz using Digidata 1550 (Molecular Devices), and recorded using pCLAMP 10.7 software (Axon Instruments). Borosilicate glass micropipettes with a tip resistance of ∼4-5 MΩ were prepared using a P-1000 puller (Sutter Instruments) and then filled with an intracellular recording solution containing (in mM) 120 gluconic acid, 15 CsCl, 10 BAPTA, 10 HEPES, 3 MgCl_2_, 1 CaCl_2_, and 2 ATP-Mg salts (pH 7.2 adjusted with CsOH). The standard extracellular recording solution (ECS) contained (in mM) 160 NaCl, 2.5 KCl, 10 HEPES, 10 D-glucose, 0.2 EDTA, and 0.7 CaCl_2_ (pH 7.3 adjusted with NaOH). ECS containing the indicated concentrations of L-glutamate and glycine were applied using a multi-barrel rapid application system with a time constant of solution exchange around the recorded cell of ∼20 ms (Skrenkova et al., 2020). All electrophysiological recordings were acquired at room temperature.

The concentration-dependent effects of agonists were analyzed using *Equation (1)*: I = 1/(1 + (EC_50_/[agonist])*^h^*), where I_max_ is the maximal steady-state current amplitude in response to agonist, EC_50_ is the concentration of the agonist eliciting half of the maximal response, [agonist] is the concentration of agonist, and h is the apparent Hill coefficient.

### 2.5. In silico experiments using molecular modeling in tandem with molecular dynamic simulations

The crystal structure of LBDs of human GluN1/GluN2A receptor in complex with GNE6901 was obtained from the RCSB PDB database (PDB ID: 5KCJ) (Hackos et al., 2016); the allosteric inhibitor was first removed from the receptor-ligand complex. The DockPrep function of UCSF Chimera software (version 1.15rc) was employed for structure preparation (Pettersen et al., 2004). Selected point mutations in the GluN1/GluN2A receptor structure were generated using the same software, utilizing the Dunbrack 2010 rotamer library (Shapovalov and Dunbrack, 2011). Gromacs version 2020.4 carried out molecular dynamic (MD) simulation. The receptor-ligand complex was solvated in the periodic water box using the SPC216 model. The system was neutralized by adding Cl^−^ and Na^+^ ions to a concentration of 10 nM. The system energy was minimized and equilibrated in a 100-ps isothermal-isochoric NVT and then a 100-ps isothermal-isobaric NPT phase. Then, a 50-ns MD simulation was run at 300 K.

### Statistical analyses

For microscopical data, statistical analysis was performed for normalized and log-transformed data to stabilize variability against the mean. Data were cleaned of outliers outside the 1.5 times interquartile range and tested for normality using the D’Agostino-Pearson test. Statistical significance was tested using a one-factor or two-factor (experiments with ARIAD-mNEON-GluN1-1a construct) ANOVA, followed by Tukey’s variant of multiple comparisons via Matlab (Matlab, 2022b; using the functions anova1, anovan, and multcompare). The rubustfit function in Matlab was used to analyze the linear regression; points whose linear regression residuals were ≥ 2 times the median absolute deviation (MAD) were identified as outliers. The regression quality was expressed as R^2^ (coefficient of determination) by calculating the 95% confidence intervals of the regression line using the predint function (parameter functional = on). Correlation strength was determined by R^2^ values, with the following classification: very strong (1.00-0.80), strong (0.79-0.60), moderate (0.59-0.40), weak (0.39-0.20), very weak (0.19-0.01), and no correlation (0.00). Differences with a p-value <0.050 were considered statistically significant; data are presented as mean ± standard error of the mean (SEM).

## Results

### Alanine substitutions of the amino acid residues of LBDs involved in direct interactions with agonists differentially alter the early transport of GluN1/GluN2A receptors

We aimed to determine whether structural changes in the LBD of the GluN1 subunit, induced by targeted substitution of amino acid residues that directly interact with its agonist glycine, alter the early trafficking of GluN1/GluN2A receptors. Previous structural studies have shown that within the LBD of the GluN1 subunit, the carboxyl group of glycine interacts with the guanidine group of residue R523, the amino groups of residues T518 and S688, and the hydroxy group of residue S688. The amino group of glycine interacts with residues P516, T518, and D732, the side chain of residue F484 is responsible for the hydrophobic interaction with glycine, and the LBD structure is formed by an interdomain interaction between residues Q405, W731, and D732 (Figure 1B) (Furukawa and Gouaux, 2003; Inanobe et al., 2005). We first decided to co-express the WT YFP-GluN1-1a or mutant YFP-GluN1-1a subunits together with the WT GluN2A subunit in HEK293 cells because (i) the GluN1-1a splice variant is retained in the ER in the absence of GluN2 subunits (Huh and Wenthold, 1999), (ii) the GluN2A subunit is the most abundantly represented GluN2 subtype in the adult forebrain (Monyer et al., 1994; Sheng et al., 1994; Maier et al., 2007), (iii) the role of LBDs of both the GluN1 or GluN2A subunits in the regulation of early transport of the GluN1/GluN2A receptor has not been comprehensively tested, and (iv) HEK293 cells do not express endogenous GluN subunits and are commonly used to study the trafficking and functional properties of the NMDARs (Collett and Collingridge, 2004). To minimize the risk of excitotoxic damage caused by the expression of NMDARs, we cultured transfected HEK293 cells in a medium supplemented with 1% FBS, 2 mM Mg^2+^, and 10 µM D-APV or 10 µM DCKA (as described in Methods). In addition, we used HEK293 cells only up to 10 passages, and each experiment had negative (i.e., YFP-GluN1-1a subunit) and positive (i.e., YFP-GluN1-1a/GluN2A receptor) controls. To accurately detect differences between WT and mutant GluN1/GluN2A receptors, we labeled surface NMDARs by combining a primary polyclonal rabbit anti-GFP antibody with a secondary Alexa 555-labeled anti-rabbit antibody. Although this combination of primary and secondary antibodies can cluster surface NMDARs, we do not foresee a bias in our microscopic data because this approach (i) provided comparable results to using fluorescently labeled anti-GFP nanobodies in our recent study (Kolcheva et al., 2023) but (ii) amplifies the surface signal of NMDARs compared to anti-GFP nanobodies, allowing more accurate detection of differences in surface expression of mutant NMDARs. Initially, we individually replaced the GluN1-Q405, GluN1-F484, GluN1-P516, GluN1-T518, GluN1-R523, GluN1-S688, GluN1-W731, and GluN1-D732 residues, with alanine residues (i.e., a neutral amino acid that should not dramatically alter protein structure) in the YFP-GluN1-1a subunit. Using anti-GFP antibody immunostaining, we measured surface and total expression levels of WT and mutant YFP-GluN1-1a/GluN2A receptors; examples of images of transfected HEK293 cells are shown in Figure 1C. We found that the surface expression of YFP-GluN1-1a/GluN2A receptors carrying alanine substitutions showed the following relationship: WT>GluN1-S688A>GluN1-Q405A>GluN1-P516A>GluN1-T518A> GluN1-F484A>GluN1-W731A>GluN1-D732A>GluN1-R523A (Figure 1D). We next performed whole-cell patch-clamp recordings from HEK293 cells expressing WT and mutant YFP-GluN1-1a/GluN2A receptors to learn how alanine substitutions in LBD of the GluN1 subunit alter EC_50_ values for glycine. We elicited the current responses using ECS containing a range of appropriate glycine concentrations (≤10 mM due to potential problems with higher osmolarity) with a fixed concentration of 300 µM L-glutamate and maintaining the membrane potential at -60 mV (Figure 1E). Our electrophysiological experiments showed that HEK293 cells expressing the YFP-GluN1-1a-D732A/GluN2A receptors did not generate current responses at any of the glycine concentrations tested, consistently with our microscopical data showing their disrupted surface expression (see Figure 1D). HEK293 cells expressing the remaining mutant YFP-GluN1-1a/GluN2A receptors produced measurable current responses induced by L-glutamate and various glycine concentrations, allowing us to generate concentration-response curves for glycine (Figure 1F). The calculated glycine EC_50_ values for YFP-GluN1-1a/GluN2A receptors carrying alanine substitutions showed the following relationship: GluN1-Q405A< WT< GluN1-S688A< GluN1-P516A< GluN1-T518A< GluN1-W731A< GluN1-F484A< GluN1-R523A (Table 1), and they were negatively strongly correlated (R^2^ = 0.70) with their surface expression (Figure 1G). Thus, our experiments showed that structural changes in LBD of the GluN1 subunit induced by alanine replacements are reflected in the surface expression of YFP-GluN1-1a/GluN2A receptors.

The CTD splice variants of the GluN1 subunit show remarkable differences in their surface expression. As described above, the GluN1-1a subunit requires the presence of the GluN2 subunit for its release from the ER; in contrast, the GluN1-4a subunit shows robust expression on the cell surface even without the GluN2 subunit (Okabe et al., 1999). Next, we performed an analogous microscopic and electrophysiological analysis in HEK293 cells transfected with WT or untagged GluN1-4a subunits carrying the alanine replacements and the WT GFP-GluN2A subunit as above with the YFP-GluN1-1a/GluN2A receptors. We found the following relationship for the relative surface expression of mutant GluN1-4a/GFP-GluN2A receptors: WT>GluN1-S688A> GluN1-Q405A> GluN1-P516A> GluN1-T518A> GluN1-F484A> GluN1-W731A> GluN1-D732A> GluN1-R523A (Figure 1H), which was negatively strongly correlated (R^2^ = 0.64) with the obtained EC_50_ values for glycine (Figure 1I, J; summarized in Table 1). In the case of HEK293 cells transfected with GluN1-4a-D732A/GFP-GluN2A receptors, we detected only the current responses <100 pA, preventing their detailed concentration-dependent analysis. The measured EC_50_ values for glycine did not differ between the corresponding mutant YFP-GluN1-1a/GluN2A and GluN1-4a/GFP-GluN2A receptors, showing that the presence of GFP or YFP tags as well as differently spliced CTDs of the GluN subunit, do not affect the EC_50_ values for glycine (Figure 1F, I). Our experiments showed that specific alanine substitutions in the LBD of the GluN1 subunit differentially alter the surface trafficking of GluN1/GluN2A receptors, independent of the splice variant of the GluN1 subunit.

We next sought to answer whether structural changes in LBD of the GluN2A subunit induced by alanine substitutions of the amino acid residues involved in its interaction with L-glutamate affect the surface expression of GluN1/GluN2A receptors. Previous structural studies with LBD of the GluN2A subunit have shown that the α-carboxyl group of L-glutamate interacts with the side group of residue R518 and with the amino groups of residues S689 and T513 and that the amino group of L-glutamate interacts with the hydroxy groups of residues S511 and T513 and, via the water molecule, with the γ-carboxyl group of residue E413 and the hydroxy group of residue Y761. Also, the γ-carboxyl group of L-glutamate interacts with the amino groups of residues S689 and T690 and the hydroxy group of residue T690. In addition, L-glutamate in LBD is stabilized by hydrophobic interactions with residues H485 and Y730, and the side chains of residues E413 and Y730 form inter-domain interactions (Figure 2A) (Laube et al., 2004; Furukawa et al., 2005; Chen and Wyllie, 2006; Maier et al., 2007; Jespersen et al., 2014). To reduce the number of the studied mutant NMDARs, the GluN2A-V685, GluN2A-G688, and GluN2A-E691 residues involved in the interaction with L-glutamate via two water molecules were not further tested (Jespersen et al., 2014). Furthermore, the GluN2A-D731 residue was omitted due to conflicting findings in previous studies, which presented varying perspectives on its interaction with L-glutamate, suggesting direct interaction (Jespersen et al., 2014), interaction via water molecules (Hansen et al., 2021), and no interaction with L-glutamate (Furukawa et al., 2005). We co-transfected HEK293 cells with WT or mutant GFP-GluN2A subunits containing the E413A, H485A, S511A, T513A, R518A, S689A, T690A, Y730A, and Y761A mutations, together with the WT GluN1-4a subunit; as this GluN subunit combination allows more straightforward electrophysiological characterization of mutant NMDARs with reduced surface expression (as shown in Figure 1). We immunofluorescently labeled HEK293 cells expressing WT and mutant GluN1-4a/GFP-GluN2A receptors using an anti-GFP antibody (Figure 2B); our subsequent microscopical analysis showed the following trend in their surface expression: GluN2A-S511A>WT> GluN2A-S689A>GluN2A-T690A> GluN2A-H485A> GluN2A-T513A> GluN2A-R518A> GluN2A-E413A> GluN2A-Y730A> GluN2A-Y761A (Figure 3C). Then, we elicited the current responses by application of L-glutamate in the appropriate concentration range (10 mM) in the continuous presence of 100 µM glycine (Figure 2D). The analysis of the concentration-response curves for L-glutamate at GluN1-4a/GFP-GluN2A receptors showed the following trend for the EC_50_ values: WT< GluN2A-S689A< GluN2A-S511A< GluN2A-T513A< GluN2A-E413A< GluN2A-Y730A<GluN2A-Y761A< GluN2A-H485A< GluN2A-T690A< GluN2A-R518A (Figure 3E and Table 2), which were negatively weakly correlated with the relative surface expression values (R^2^ = 0.39), except for a GluN2A-S511A mutation (Figure 2F). These results also showed that the structural changes in LBD of the GluN2A subunit induced by alanine replacements are reflected in the regulation of the surface expression of the GluN1/GluN2A receptors.

**Figure 2.**
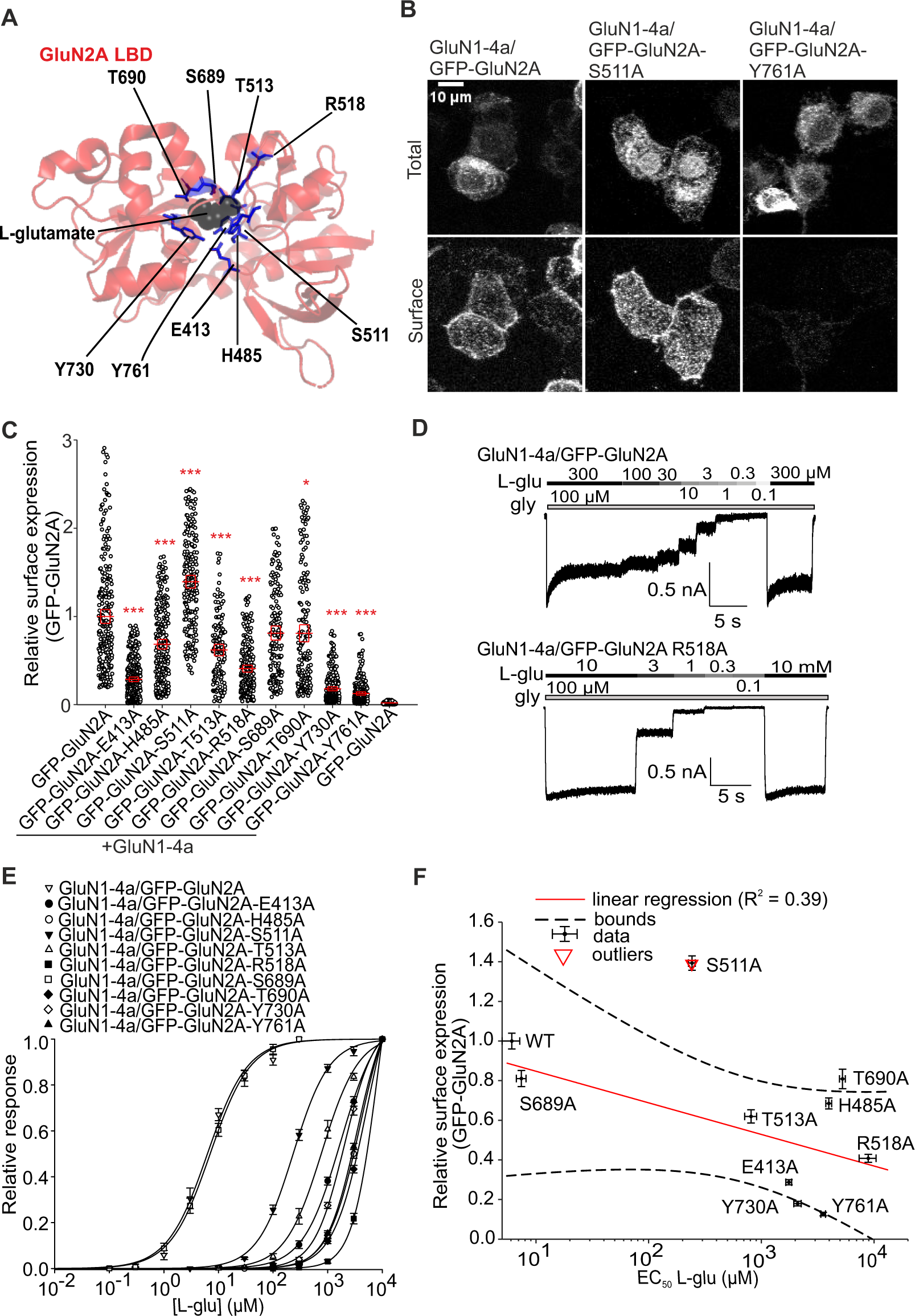
Mutations of residues in LBD of GluN2A subunit interacting with L-glutamate affect surface expression and L-glutamate potency of GluN1/GluN2A receptors. (**A**) Structural model of the LBD of the GluN2A subunit labeled with L-glutamate-interacting amino acids (based on PDB ID: 5I57). (**B**) Representative images of HEK293 cells co-transfected with GluN1-4a and WT or mutated GFP-GluN2A subunits. The total and surface signals (top and bottom row, respectively) of GFP-GluN2A subunits were labeled using an anti-GFP antibody 24 hours after the transfection. (**C**) Summary of the relative surface expression of NMDARs consisting of either WT or mutated GFP-GluN2A subunit together with GluN1-4a subunit, measured using fluorescence microscopy; \**p* < 0.050 and \*\*\**p* < 0.001 vs WT; one-way ANOVA. Data points correspond to individual cells (*n* ≥ 136), and the red box plot represents mean ± SEM. (**D**) Representative whole-cell voltage-clamp recordings of HEK293 cells transfected with the indicated NMDAR subunits. L-glutamate (L-glu) at the indicated concentrations was applied in the continuous presence of 100 µM glycine (gly). (**E**) Normalized concentration-response curves for L-glutamate measured from HEK293 cells expressing NMDARs containing WT or mutated GluN1-4a/GFP-GluN2A receptors were obtained by fitting the data using Equation (1) (see Methods) for the summary of fitting parameters see **Table 2**. (**F**) Correlation of surface expression and L-glutamate EC_50_ values at NMDARs composed of WT or mutated GluN1-4a/GFP-GluN2A receptors; data were fitted by linear regression and not correlating mutation (outlier) is labeled in red.

**Figure 3.**
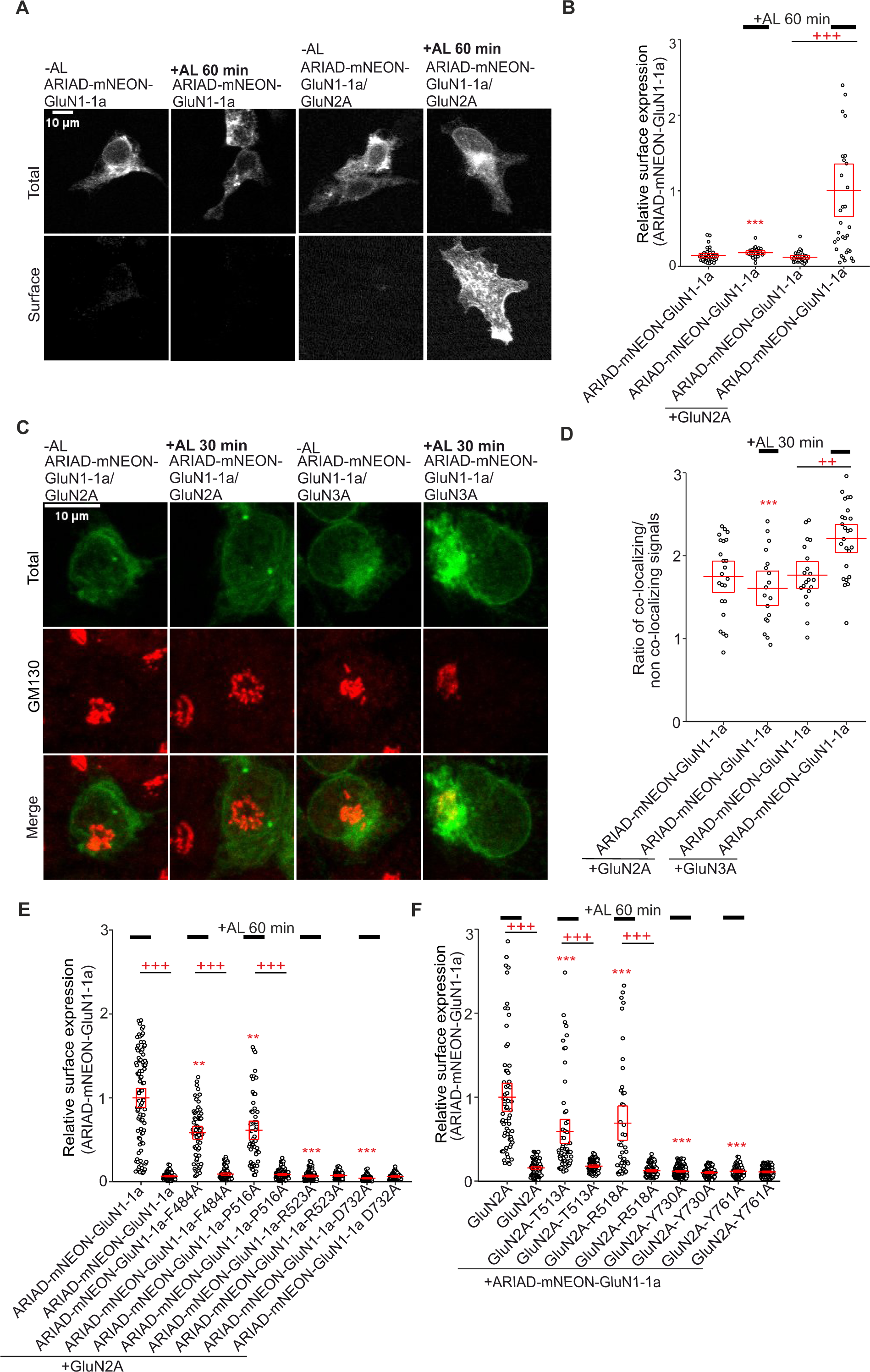
Alanine substitutions in the LBDs of both GluN1 and GluN2A subunits alter the early trafficking of NMDARs. **(A)** Representative images of HEK293 cells expressing ARIAD-mNEON-GluN1-1a construct either alone or with the GluN2A subunit at 0 min (without AL) and 60 min after adding AL. The total and surface signals (top and bottom row, respectively) of ARIAD-mNEON-GluN1-1a subunits were labeled using anti-mNEONGreen antibody 24 hours after the transfection. The representative images of GluN1-4a/ARIAD-mNEON-GluN2A receptors are shown in Figure 3-1. **(B)** Summary of relative surface expression of NMDARs containing WT or mutated ARIAD-mNEON-GluN1-1a construct together with GluN2A subunit in the absence or presence (60 min) of AL, measured using fluorescence microscopy; \*\*\**p*< 0.001 for differences between ARIAD-mNEON-GluN1-1a/GluN2A in presence of AL and mutated NMDARs; +++*p* < 0.001 for differences between presence and absence of AL; two-way ANOVA. Data points correspond to individual cells (*n* ≥ 26), and the red box plot represents mean ± SEM. See Figure 3-2A and Figure 3-2B for a summary of the surface expression of the time points for ARIAD-mNEON-GluN1-1a with GluN2A or GluN3A subunits. (**C**) Representative confocal microscopy images of the HEK293 cells co-transfected with ARIAD-mNEON-GluN1-1a construct and GluN2A or GluN3A subunits in the absence or presence (30 min) of AL; the anti-GM130 antibody was used to label the GA. (**D**) Summary of the average intensity of ARIAD-mNEON-GluN1-1a subunit signal colocalized with GM130 over the average intensity of ARIAD-mNEON-GluN1-1a subunit signal outside the GM130 signal, calculated for the indicated NMDAR combinations; \*\*\**p* < 0.001 for differences between ARIAD-mNEON-GluN1-1a/GluN3A receptors in presence of AL and mutated NMDARs; ++*p* < 0.01 for differences between presence and absence of AL; two-way ANOVA. Data points correspond to individual cells (*n* ≥ 18), and the red box plot represents mean ± SEM. See Figure 3-3 for a summary of the time points for colocalized ARIAD-mNEON-GluN1-1a with GluN2A subunits. (**E, F**) Summary of relative surface expression of NMDARs containing WT or mutated ARIAD-mNEON-GluN1-1a subunit co-expressed together with WT GluN2A subunit **(E)** or WT ARIAD-mNEON-GluN1-1a subunit co-expressed with WT or mutated GluN2A subunit **(F)** measured in the absence or presence (60 min) of AL, measured using fluorescence microscopy; \*\**p* < 0.010, and \*\*\**p* < 0.001 for differences between ARIAD-mNEON-GluN1-1a/GluN2A receptor and mutated NMDARs in presence of AL; +++*p* < 0.001 for differences between presence and absence of AL; two-way ANOVA. Data points correspond to individual cells (*n* ≥ 55 for **E** and *n* ≥ 60 for **F**), and the red box plot represents mean ± SEM.

**Table 2.** Summary of fitting parameters for steady-state concentration-response curves for L-glutamate effect measured in HEK293 cells expressing the indicated NMDAR subunits carrying a mutation in the LBD of GluN2A subunit.

| **Receptor** | **EC<sub>50</sub> (μM)** | ***h*** | ***n*** |
| --- | --- | --- | --- |
| **GluN1-4a/**<br>**GFP-GluN2A** | 6.10 ± 0.70 | 1.34 ± 0.04 | 5 |
| **GluN1-4a/**<br>**GFP-GluN2A-E413A** | 1,754.81 ± 114.61 | 1.27 ± 0.05 | 6 |
| **GluN1-4a/**<br>**GFP-GluN2A-H485A** | 3,969.98 ± 253.57 | 1.50 ± 0.04 | 6 |
| **GluN1-4a/**<br>**GFP-GluN2A-S511A** | 235.41 ± 18.32 | 1.35 ± 0.03 | 6 |
| **GluN1-4a/**<br>**GFP-GluN2A-T513A** | 809.56 ± 109.74 | 1.33 ± 0.03 | 5 |
| **GluN1-4a/**<br>**GFP-GluN2A-R518A** | 8,895.17 ± 1,702.60 | 1.96 ± 0.15 | 5 |
| **GluN1-4a/**<br>**GFP-GluN2A-S689A** | 7.42 ± 0.82 | 1.17 ± 0.05 | 5 |
| **GluN1-4a/**<br>**GFP-GluN2A-T690A** | 5,254.10 ± 339.72 | 1.45 ± 0.05 | 7 |
| **GluN1-4a/**<br>**GFP-GluN2A-Y730A** | 2,099.49 ± 170.86 | 1.58 ± 0.07 | 6 |
| **GluN1-4a/**<br>**GFP-GluN2A-Y761A** | 3,542.66 ± 173.32 | 1.58 ± 0.10 | 8 |

The EC<sub>50</sub> and *h* values were obtained by fitting the data from individual cells to Equation (1).

We further used the ARIAD system, which allows synchronized release of membrane proteins from the ER by the addition of the ARIAD ligand (AL) (Hangen et al., 2018), to test the hypothesis that changes in surface expression of mutant GluN1/GluN2A receptors are caused at the level of early transport. We created ARIAD-mNEON-GluN1-1a and ARIAD-mNEON-GluN2A constructs and expressed them separately in HEK293 cells. Unfortunately, HEK293 cells expressing the ARIAD-mNEON-GluN2A construct showed a distinct surface signal detected by the anti-mNEONGreen antibody 24 hours after transfection, even without adding AL (0 min, Figure 3-1). In contrast, HEK293 cells expressing the ARIAD-mNEON-GluN1-1a construct showed no surface signal either under control conditions (0 min) or 60 min after addition of AL (Figure 3A). Co-transfection of HEK293 cells with the ARIAD-mNEON-GluN1-1a construct and the WT GluN2A subunit resulted in robust surface expression of the mNEON-GluN1-1a/GluN2A receptors 60 min after addition of AL but not under control conditions (0 min), confirming the functionality of the ARIAD-mNEON-GluN1-1a construct for studying early transport of GluN1/GluN2A receptors (Figure 3B; time points 0, 30, 60 and 120 min after the addition of AL shown in Figure 3-2A and for ARIAD-mNEON-GluN1/GluN3A in Figure 3-2B). Given that only a smaller fraction of functional GluN1/GluN2A receptors likely leave the ER structures in HEK293 cells (Huh and Wenthold, 1999; Hawkins et al., 2004), their co-localization with GA is used as a measure to quantify the early transport of NMDAR (She et al., 2012; Skrenkova et al., 2019). Therefore, we co-transfected HEK293 cells with the ARIAD-mNEON-GluN1-1a construct, in combination with the WT GluN2A subunit (or with the WT GluN3A subunit as a positive control) and labeled GA structures on fixed cells with anti-GM130 antibody at 0 and 30 min after addition of AL (Figure 3C; time points 0, 30, 60 and 120 min after the addition of AL shown in Figure 3-3). Consistent with our previous experiments with GluN1/GFP-GluN3A receptors (Skrenkova et al., 2019), our microscopical analysis showed that mNEON-GluN1-1a/GluN3A receptors exhibited a higher rate of co-localization with GA structures 30 min after addition of AL compared with negative control (0 min). In contrast, mNEON-GluN1-1a/GluN2A receptors did not show a different rate of co-localization with GA structures 30 min after the addition of AL compared with negative control (0 min, Figure 3D); we obtained the same conclusion in another experiment involving time points 0, 15, 30, 45 and 60 minutes after the addition of AL (Figure 3-3). One possible explanation is that GluN1/GluN2A receptors can bypass GA via an unconventional pathway during early transport to the cell surface (see Discussion). Therefore, we did not further analyze the co-localization of mutant GluN1/GluN2A receptors with GA structures to measure their early transport, and we focused on characterizing their surface expression 60 min after adding AL.

To test whether mutant GluN1/GluN2A receptors exhibit altered early transport, we introduced four alanine substitutions into the ARIAD-mNEON-GluN1-1a construct, which caused the most significant reductions (∼95% GluN1-R523A and ∼90% GluN1-D732A) and moderate reductions (∼53% GluN1-F484A and ∼43% GluN1-P516A) in surface expression of GluN1/GluN2A receptors (Figure 1). Consistent with our previous results, GluN1-R523A and GluN1-D732A mutations eliminated surface expression (∼7% for GluN1-R523 and ∼4% for GluN1-D732A). In contrast, both GluN1-F484A and GluN1-P516A mutations caused ∼42% resp. ∼39% reduction in surface expression of mNEON-GluN1-1a/GluN2A receptors measured 60 min after addition of AL (Figure 3E). For further analysis using the ARIAD system, we selected, based on previous experiments, four alanine substitutions in the LBD of the GluN2A subunit, two that caused the most significant reduction (∼82% GluN2A-Y730A and ∼87% GluN2A-Y761A) and two that showed moderate reduction (∼38% GluN2A-T513A and ∼59% GluN2A-R518A) in surface expression of GluN1/GluN2A receptors (Figure 2). Our analysis confirmed that the GluN2A-Y730A and GluN2A-Y761A mutations were essentially eliminated (both ∼12%) and that the GluN2A-T513A and GluN2A-R518A mutations caused a ∼41% resp. ∼31% reduction in surface expression of mNEON-GluN1-1a/GluN2A receptors in HEK293 cells at 60 min after addition of AL (Figure 3F). Therefore, we suggest that the observed decrease in surface expression of GluN1/GluN2A receptors with alanine substitutions in the LBDs of both GluN1 and GluN2A subunits is due to their altered early transport, likely during their processing in the ER.

We then investigated whether structural changes in LBDs of GluN1 and GluN2A subunits induced by alanine substitutions are sensed independently by control mechanisms. Therefore, we co-transfected HEK293 cells with a combination of WT and mutant GluN1-4a (containing F484A or P516A mutations) and GFP-GluN2A (containing T513A or R518A mutations) subunits. Our analysis showed that in all cases, the combination of two alanine substitutions (GluN1-4a-F484A/GFP-GluN2A-T513A, GluN1-4a-F484A/GFP-GluN2A-R518A, GluN1-4a-P516A/GFP-GluN2A-T513A, and GluN1-4a-P516A/GFP-

GluN2A-R518A) caused significantly reduced surface expression of the GluN1-4a/GFP-GluN2A receptors (∼90%) compared to the GluN1-4a/GFP-GluN2A receptors containing only single alanine substitutions (Figure 4A). Thus, our experiments showed that structural changes in the LBDs of GluN1 and GluN2A subunits additively regulate the early transport of GluN1/GluN2A receptors and are likely recognized independently by control mechanisms.

**Figure 4.**
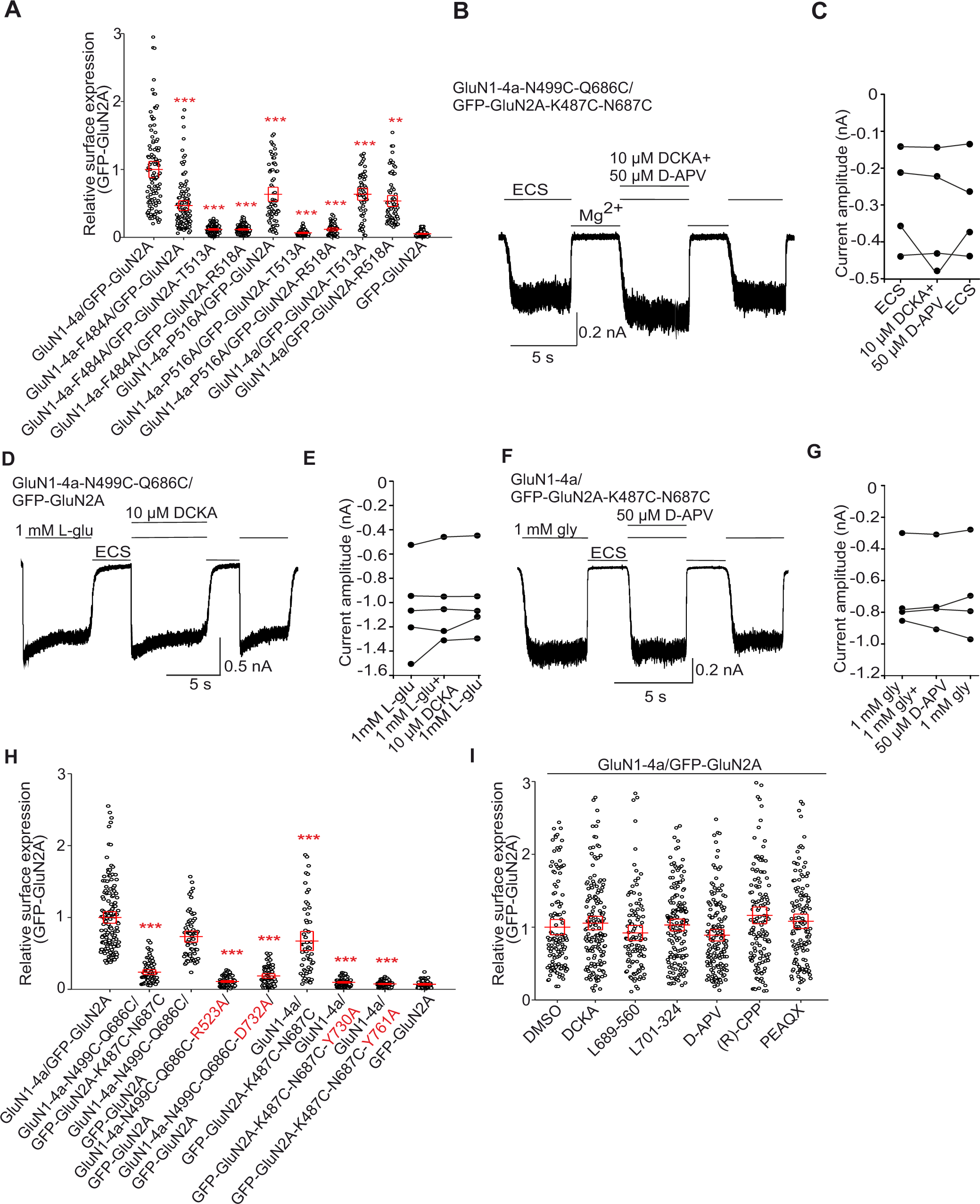
Structural changes in the LBDs of GluN1 and GluN2A subunits additively regulate the early trafficking of GluN1/GluN2A receptors. (**A**) Summary of the relative surface expression of NMDARs containing mutations within LBDs of both GluN1-4a and GFP-GluN2A subunits, measured using fluorescence microscopy; \*\**p* < 0.01, and \*\*\**p* < 0.001 vs WT; one-way ANOVA. Data points correspond to individual cells (*n* ≥ 55), and the red box plot represents mean ± SEM. (**B,D,F**) Representative whole-cell recordings of GluN1-4a-N499C-Q686C/GFP-GluN2A-K487C-N687C (**B**), GluN1-4a-N499C-Q686C/GFP-GluN2A (**D**), and GluN1-4a/GFP-GluN2A-K487C-N687C receptors (**F**) expressed in the HEK293 cells. The current responses were elicited under the specified conditions. ECS denotes Mg^2+^-free extracellular solution, L-glu represents L-glutamate (1 mM), and gly indicates glycine (1 mM). As indicated, 10 µM 5,7-dichlorokynurenic acid (DCKA) and 50 µM D-2-amino-5-phosphonopentanoate (D-APV) were applied. (**C,E,G**) Comparison of current response amplitudes evoked by ECS without Mg^2+^ (**C**), 1 mM L-glutamate plus 10 µM DCKA **(E)**, and 1 mM glycine plus 50 µM D-APV **(G)**. (**H**) Summary of relative surface expression for NMDARs composed of GluN1-4a and GFP-GluN2A subunits with the closed cleft conformation of LBDs in combination with selected alanine substitutions, measured using fluorescence microscopy; \*\*\**p* < 0.001 vs WT; one-way ANOVA. Data points correspond to individual cells (*n* ≥ 54), and the red box plot represents mean ± SEM. (**I**) Summary of the effect of indicated glycine or L-glutamate binding sites competitive antagonists on surface expression of GluN1-4a/GFP-GluN2A receptors, measured using fluorescence microscopy; no significance; one-way ANOVA. Their respective abbreviations are as follows: DCKA, 5,7-dichlorokynurenic acid; L-689,560, 4-trans-2-carboxy-5,7-dichloro-4-phenylaminocarbonylamino-1,2,3,4-tetrahydroquinoline; L-701,324, 7-chloro-4-hydroxy-3-(3-phenoxy)phenyl-2(1H)-quinolone; D-APV, D-2-Amino-5-phosphonopentanoate, PEAQX, 5-phosphonomethyl-1,4-dihydroquinoxaline-2,3-dione; CPP, 4-(3-phosphonopropyl) pizerazine-2-carboxylic acid. Data points correspond to individual cells (*n* ≥ 107), and the red box plot represents mean ± SEM.

Previous studies have found that the introduction of specific cysteine residues into LBDs in the GluN1 (N499C-Q686C) and GluN2A (K487C-N687C) subunits leads to the closed cleft conformation of the LBDs (Blanke and VanDongen, 2008; Kussius and Popescu, 2010; Dai and Zhou, 2016). We first verified by electrophysiological measurements that GluN1-4a-N499C-Q686C/GFP-GluN2A-K487C-N687C receptors expressed in HEK293 cells generate current responses elicited by ECS application in the absence of glycine and L-glutamate; also in the presence of the competitive antagonists (10 µM DCKA and 50 µM D-APV; Figure 4B, C). Subsequently, we found that HEK293 cells expressing GluN1-4a-N499C-Q686C/GFP-GluN2A receptors produced current responses in the presence of 1 mM L-glutamate only, which were insensitive to 10 µM DCKA (Figure 4D,E). In contrast, HEK293 cells expressing GluN1-4a/GFP-GluN2A-K487C-N687C receptors had current responses elicited by 1 mM glycine only, which were insensitive to 50 µM D-APV (Figure 4F,G). Subsequent microscopical experiments showed that GluN1-4a-N499C-Q686C/GFP-GluN2A-K487C-N687C receptors exhibit profoundly reduced surface expression (∼76%), the GluN1-4a-N499C-Q686C/GFP-GluN2A receptors showed a tendency in reduction (∼27%), and GluN1-4a/GFP-GluN2A-K487C-N687C showed a significant decrease (∼33%) in surface expression compared to WT GluN1-4a/GFP-GluN2A receptors (Figure 4H). These data support the hypothesis that structural changes in the LBDs of GluN1 and GluN2A subunits additively regulate early transport of GluN1/GluN2A receptors.

We next asked whether introducing alanine substitutions in the LBDs of the GluN1 (R523A and D732A) or GluN2A (Y730A and Y761A) subunits, which caused a profound reduction in surface expression of GluN1/GluN2A receptors, also affect surface expression of GluN1/GluN2A receptors with closed cleft conformation of LBD. Our microscopical analysis showed that single alanine substitutions in the closed cleft conformation of LBDs of both GluN1 and GluN2A subunits, i.e., in the case of GluN1-4a-N499C-R523A-Q686C/GFP-GluN2A, GluN1-4a-N499C-Q686C-D732A/GFP-GluN2A, GluN1-4a/GFP-GluN2A-K487C-N687C-Y730A, and GluN1-4a/GFP-GluN2A-K487C-N687C-Y761A receptors (Figure 4H), induced a profound decrease of their surface expression, comparable to the decline observed for GluN1-4a/GFP-GluN2A receptors carrying only the corresponding alanine substitutions. These experiments demonstrated that the control mechanisms perceive structural changes induced by alanine substitutions regardless of the closed cleft conformation of LBDs of both GluN1 and GluN2A subunits.

Finally, we investigated whether the presence of selected competitive antagonists affects the surface expression of GluN1/GluN2A receptors. Specifically, we used the commonly used DCKA and two highly potent antagonists, L689-560 and L701-324, acting via the GluN1 subunit, and the widely used D-APV and two highly potent antagonists, (R)-CPP and PEAQX, with high selectivity for the GluN2A subunit (Traynelis et al., 2010). We cultured the HEK293 cells expressing WT GluN1-4a/GFP-GluN2A receptors for 36 hours after the completion of the transfection in the presence of 300 µM of competitive antagonists, as higher concentrations could mediate toxic effects on HEK293 cells (data not shown). We did not observe any differences in the surface expression level of WT GluN1-4a/GFP-GluN2A receptors in the presence of any competitive antagonists (Figure 4I), indicating that the control mechanisms do not perceive structural changes induced by the interaction of the competitive antagonists with the GluN1/GluN2A receptors. However, we cannot exclude that our observation is due to the limited presence of competitive antagonists in intracellular compartments, including the ER.

### Pathogenic variants and the mutant cycles of the specific amino acid residues in LBDs of GluN1 and GluN2A subunits indicate tight regulation of early trafficking of GluN1/GluN2A receptors

We have searched the "GRIN variants DATABASE" (available at URL: https://alf06.uab.es/grindb/home) for pathogenic and potentially pathogenic "missense" variants (hereafter referred to as "pathogenic variants") of amino acid residues in LBDs involved in the direct interaction of glycine with the GluN1 subunit and L-glutamate with the GluN2A subunit; their list as of January 1, 2022, including relevant information, is provided in Table 3. To examine the effect of the pathogenic variants on the surface expression of GluN1/GluN2A receptors, we chose to combine YFP-hGluN1-1a and hGluN2A subunits, given that we do not have a functional tagged version of the hGluN2A subunit. We labeled the HEK293 cells transfected with WT or mutated YFP-hGluN1-1a/hGluN2A subunit using an anti-GFP antibody (Figure 5A); analysis of microscopical data revealed the following trends of surface expression levels for GluN1 subunit: GluN1-S688Y> WT> GluN1-D732E> GluN1-S688P> GluN1-R523C (Figure 5B), and for GluN2A: WT> GluN2A-T513I> GluN2A-T690M> GluN2A-S511L> GluN2A-R518L> GluN2A-R518C> GluN2A-R518H (Figure 5C) subunits. We then performed the same electrophysiological approach described above using the HEK293 cells expressing WT and mutant YFP-hGluN1-1a/hGluN2A receptors, and we analyzed concentration-response curves for glycine (for pathogenic variants in the GluN1 subunit) and L-glutamate (for pathogenic variants in the GluN2A subunit) (Figure 5D). Concerning the pathogenic variants in the GluN1 subunit, we did not detect any current responses in HEK293 cells transfected with YFP-hGluN1-1a-R523C/hGluN2A receptor. In contrast, we observed a profound 3,188-fold increase in EC_50_ values for glycine with YFP-hGluN1-1a-S688P/hGluN2A, a 944-fold increase for YFP-hGluN1-1a-S688Y/hGluN2A, and a 1,457-fold increase for YFP-hGluN1-1a-D732E/hGluN2A (Figure 5E).

**Figure 5.**
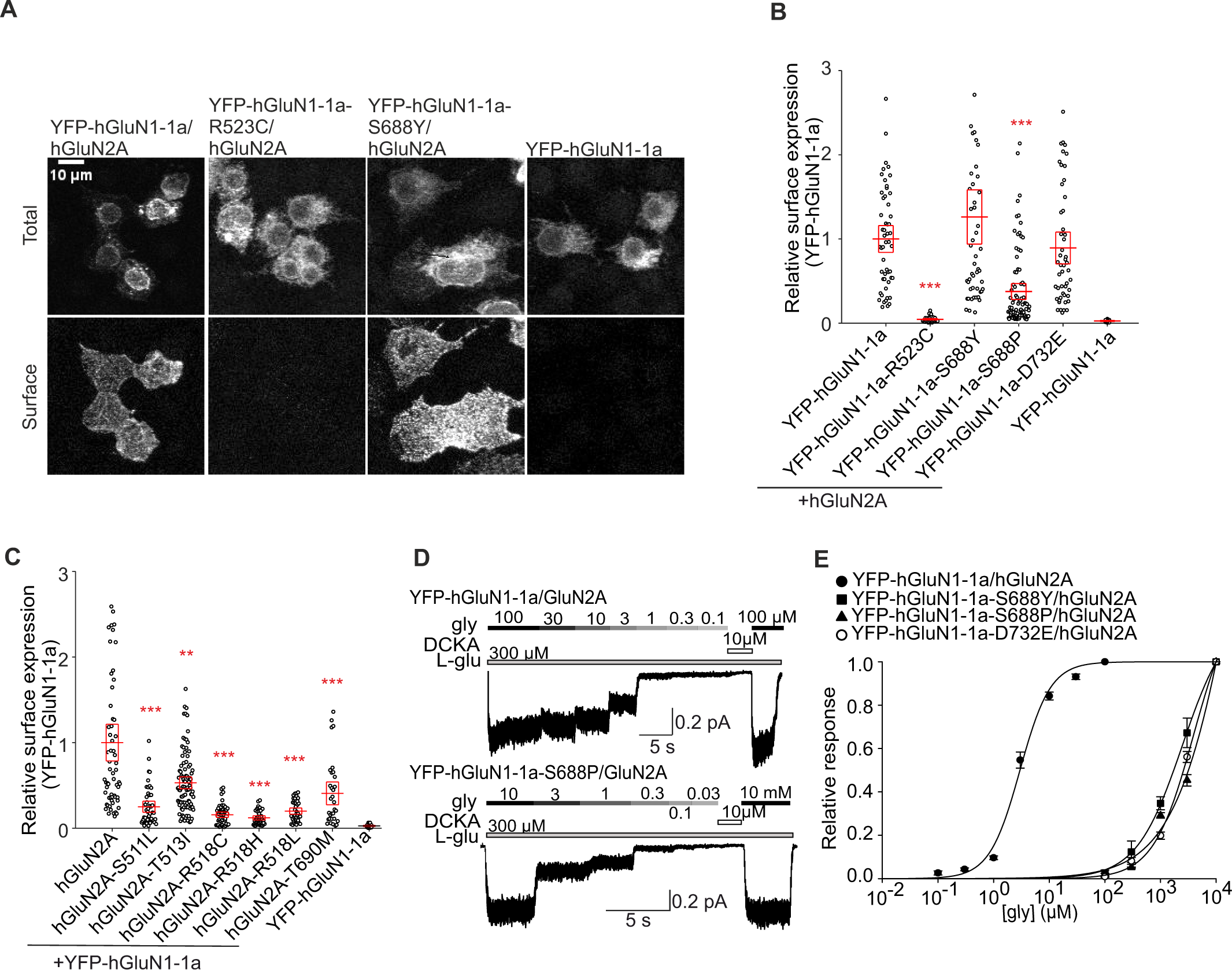
Pathogenic variants in LBDs of both GluN1 and GluN2A subunits affect the surface expression and agonist potency of NMDARs. (**A**) Representative images from HEK293 cells transfected with the WT or mutated YFP-hGluN1-1a subunit either alone or with the hGluN2A subunit. The total and the surface signals (top and bottom row, respectively) of YFP-hGluN1-1a subunits were labeled using an anti-GFP antibody 24 hours after the transfection. (**B,C**) Summary of relative surface expression of YFP-hGluN1-1a/hGluN2A receptors containing pathogenic variants within the LBDs of the YFP-hGluN1-1a subunit **(B)** or hGluN2A subunit **(C)**, measured using fluorescence microscopy; \*\**p* < 0.010, \*\*\**p* < 0.001 vs respective WT; one-way ANOVA. Data points correspond to individual cells (*n* ≥ 16 for **B**, *n* ≥ 25 for **C**), and the red box plot represents mean ± SEM. (**D**) Representative whole-cell voltage-clamp recordings from HEK293 cells transfected with the indicated NMDAR subunits. Glycine (gly) at the indicated concentrations was applied in the continuous presence of 300 µM L-glutamate (L-glu). (**E**) Normalized concentration-response curves for glycine measured from HEK293 cells expressing WT or mutated YFP-hGluN1-1a/hGluN2A receptors obtained by fitting the data using Equation (1) (see Methods); for the summary of fitting parameters, see **Table 4**.

**Table 3.** Pathogenic and potential pathogenic variants in LBDs of GluN1 and GluN2A investigated in this study.

| **Variant** | **Gene** | **Genotype** | **Phenotype** | **References** |
| --- | --- | --- | --- | --- |
| **GluN1-R523C** | *GRIN1* | c.1567C>T | DA | (Kalbaugh et al., 2004); ClinVar ID: 1308763 |
| **GluN1-S688Y** | *GRIN1* | c.2063C>A | Early infantile encephalopathy, Intellectual disability | (Zehavi et al., 2017; Skrenkova et al., 2020; Chen et al., 2023); ClinVar ID: 981272 |
| **GluN1-S688P** | *GRIN1* | c.2062T>C | DA | ClinVar ID: 429761 |
| **GluN1-D732E** | *GRIN1* | - | DA | (Santos-Gómez et al., 2021) |
| **GluN2A-S511L** | *GRIN2A* | c.1532C>T | Landau-Kleffner syndrome | ClinVar ID: 80290 |
| **GluN2A-T513I** | *GRIN2A* | c.1538C>T | Landau-Kleffner syndrome | ClinVar ID: 424392 |
| **GluN2A-R518C** | *GRIN2A* | c.1552C>T | Landau-Kleffner syndrome | (Strehlow et al., 2019); ClinVar ID: 205687 |
| **GluN2A-R518H** | *GRIN2A* | c.1553G>A | Landau-Kleffner syndrome | (Lesca et al., 2013; Conroy et al., 2014; Swanger et al., 2016; Strehlow et al., 2019); ClinVar ID: 88732 |
| **GluN2A-R518L** | *GRIN2A* | c.1553G>T | Landau-Kleffner syndrome | ClinVar ID: 567708 |
| **GluN2A-T690M** | *GRIN2A* | c.2069C>T | Inborn genetic diseases, Landau-Kleffner syndrome, neurodevelopmental disorder | (Nykamp et al., 2017; Fernández-Marmiesse et al., 2019); ClinVar ID: 444362 |

The variants were selected from the open-source GRIN database (https://alf06.uab.es/grindb/home). The information provided is current as of January 1, 2022. Variants marked as 'DA' denote disease-associated or potential pathogenic variants.

**Table 4.** Summary of fitting parameters for steady-state concentration-response curves for glycine and L-glutamate measured in HEK293 cells expressing the human versions of the GluN1/GluN2A receptors carrying the indicated pathogenic variants and the mutant cycle substitutions.

| Receptor | EC<sub>50</sub> glycine (μM) | *h* | *n* | Receptor | EC<sub>50</sub> L-glutamate (μM) | *h* | *n* |
| --- | --- | --- | --- | --- | --- | --- | --- |
| YFP-hGluN1-1a/ hGluN2A | 2.96 ± 0.29 | 0.96 ± 0.08 | 6 | YFP-hGluN1-1a/ hGluN2A | 5.31 ± 0.88 | 1.28 ± 0.04 | 5 |
| YFP-hGluN1-1a-R523C/ hGluN2A | NR | - | 10 | YFP-hGluN1-1a/ hGluN2A-S511L | NR | - | 11 |
| YFP-hGluN1-1a-S688A/ hGluN2A | 5.02 ± 0.75 | 0.96 ± 0.04 | 5 | YFP-hGluN1-1a/ hGluN2A-S511A | 179.45 ± 29.45 | 1.05 ± 0.02 | 5 |
| YFP-hGluN1-1a-S688Y/ hGluN2A | 2,795.62 ± 787.88 | 1.12 ± 0.11 | 5 | YFP-hGluN1-1a/ hGluN2A-S511P | 67.71 ± 8.08 | 1.14 ± 0.04 | 4 |
| YFP-hGluN1-1a-S688P/ hGluN2A | 9,435.59 ± 1,533.21 | 0.96 (Fixed) | 5 | YFP-hGluN1-1a/ hGluN2A-S511C | 264.49 ± 38.94 | 1.00 ± 0.04 | 5 |
| YFP-hGluN1-1a-S688C/ hGluN2A | 26.88 ± 5.07 | 0.97 ± 0.07 | 5 | YFP-hGluN1-1a/ hGluN2A-S511N | NR | - | 8 |
| YFP-hGluN1-1a-S688N/ hGluN2A | 147.66 ± 9.17 | 1.38 ± 0.08 | 7 | YFP-hGluN1-1a/ hGluN2A-S511E | 342.32 ± 61.05 | 1.13 ± 0.06 | 6 |
| YFP-hGluN1-1a-S688E/ hGluN2A | 3,044.67 ± 496.54 | 1.49 ± 0.15 | 4 | YFP-hGluN1-1a/ hGluN2A-S511K | NR | - | 8 |
| YFP-hGluN1-1a-S688K/ hGluN2A | 18.12 ± 1.67 | 1.25 ± 0.09 | 6 | YFP-hGluN1-1a/ hGluN2A-S511W | NR | - | 5 |
| YFP-hGluN1-1a-S688W/ hGluN2A | 2,249.31 ± 105.92 | 1.49 ± 0.04 | 8 | YFP-hGluN1-1a/ hGluN2A-T513I | NA | - | 7 |
| YFP-hGluN1-1a-D732A/ hGluN2A | NA | - | 5 | YFP-hGluN1-1a/ hGluN2A-R518C | NA | - | 5 |
| YFP-hGluN1-1a-D732P/ hGluN2A | NR | - | 10 | YFP-hGluN1-1a/ hGluN2A-R518H | NR | - | 10 |
| YFP-hGluN1-1a-D732C/ hGluN2A | 3,342.35 ± 768.26 | 1.09 ± 0.19 | 6 | YFP-hGluN1-1a/ hGluN2A-R518L | NR | - | 10 |
| YFP-hGluN1-1a-D732N/ hGluN2A | 19,979.95 ± 2,785.62 | 0.96 (fixed) | 4 | YFP-hGluN1-1a/ hGluN2A-T690A | 4,206.12 ± 292.47 | 1.00 ± 0.01 | 5 |
| YFP-hGluN1-1a-D732E/ hGluN2A | 4,314.16 ± 874.98 | 1.22 ± 0.08 | 4 | YFP-hGluN1-1a/ hGluN2A-T690P | NR | - | 6 |
| **YFP-hGluN1-1a-D732K/hGluN2A** | 2.07 ± 0.09 | 1.17 ± 0.02 | 7 | **YFP-hGluN1-1a/hGluN2A-T690C** | 539.13 ± 48.10 | 0.90 ± 0.06 | 8 |
| **YFP-hGluN1-1a-D732W/hGluN2A** | NR | - | 12 | **YFP-hGluN1-1a/hGluN2A-T690N** | NR | - | 8 |
|  |  |  |  | **YFP-hGluN1-1a/hGluN2A-T690M** | NR | - | 10 |
|  |  |  |  | **YFP-hGluN1-1a/hGluN2A-T690E** | NR | - | 8 |
|  |  |  |  | **YFP-hGluN1-1a/hGluN2A-T690K** | NR | - | 7 |
|  |  |  |  | **YFP-hGluN1-1a/hGluN2A-T690W** | NR | - | 6 |

The EC<sub>50</sub> and *h* values were obtained by fitting the data from individual cells to Equation (1). NR indicates “not responding”, and NA indicates “not analyzed”.

Concerning the pathogenic variants in the GluN2A subunit, we did not detect any current responses of YFP-hGluN1-1a/hGluN2A-S511L, YFP-hGluN1-1a/hGluN2A-R518H, YFP-hGluN1-1a/hGluN2A-R518L and YFP-hGluN1-1a/hGluN2A-T690M receptors even in the presence of 10 mM L-glutamate. In the case of the YFP-hGluN1-1a/hGluN2A-T513I and YFP-hGluN1-1a/hGluN2A-R518C receptors, we observed current responses only in the presence of 3 and 10 mM L-glutamate, and we did not proceed with the concentration-response analysis. These experiments showed that pathogenic variants in LBDs can have a prominent impact, distinct from the corresponding alanine substitutions, on regulating the surface expression of GluN1/GluN2A receptors.

Interestingly, the surface expression levels determined for YFP-hGluN1-1a/hGluN2A receptors containing pathogenic variants were profoundly different compared to the corresponding alanine substitutions for GluN1-S688 (∼76% for the GluN1-S688A mutation compared to ∼126% for the hGluN1-S688Y or ∼38% for the hGluN1-S688P variants; ∼10% for the GluN1-D732A mutation versus ∼89% for the hGluN1-D732E variant; ∼139% for the GluN2A-S511A mutation versus ∼25% for the hGluN2A-S511L variant; ∼81% for the GluN2A-T690A mutation versus ∼41% for the hGluN2A-T690M variant). Therefore, we prepared a series of mutant cycles of GluN1-S688, GluN1-D732, GluN2A-S511, and GluN2A-T690 residues to elucidate how their structural substitutions affect the surface expression of GluN1/GluN2A receptors. Specifically, we introduced substitutions in the YFP-hGluN1-1a and hGluN2A subunits for alanine (A; small uncharged), cysteine (C; nucleophilic), proline (P; hydrophobic), tryptophan (W; aromatic), glutamic acid (E; acidic), asparagine (N; amide), and lysine (K; basic). Regarding the mutant cycle of the GluN1-S688 residue, our microscopical analysis showed the following trend for the surface expression levels of YFP-hGluN1-1a/hGluN2A receptors expressed in HEK293 cells: WT>GluN1-S688N> GluN1-S688A> GluN1-S688E> GluN1-S688C> GluN1-S688W> GluN1-S688P>GluN1-S688K (Figure 6A). Our electrophysiological measurements showed that all YFP-hGluN1-1a/hGluN2A receptors containing the mutant GluN1-S688 residue cycle produced sufficient amplitudes of the current responses to obtain the EC_50_ values for glycine (Table 4, Figure 6B). Subsequent analysis showed no correlation between the surface expression levels and EC_50_ values for glycine for WT and mutant YFP-hGluN1-1a/hGluN2A receptors at the GluN1-688 position (R^2^ = 0.00) (Figure 6C).

**Figure 6.**
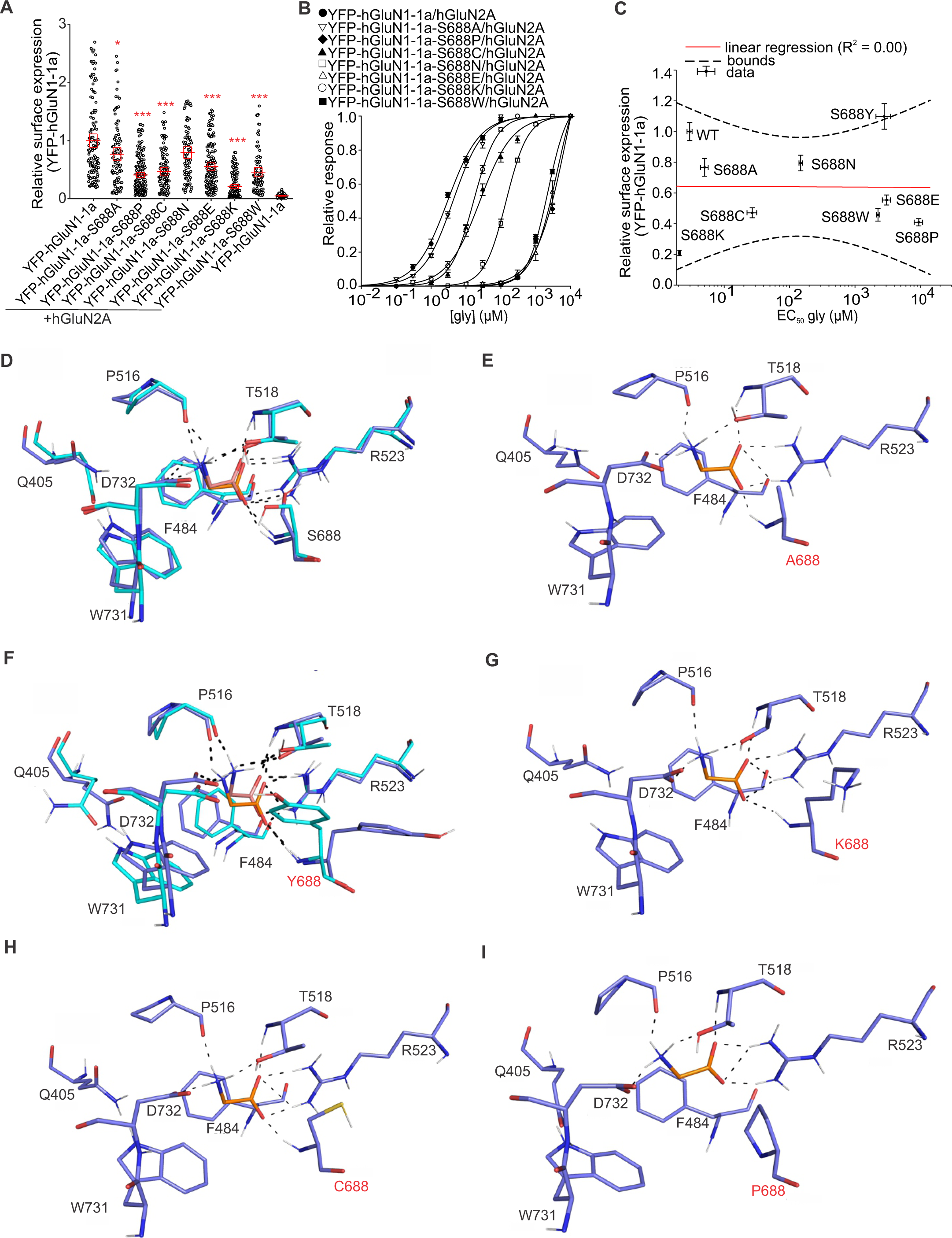
The mutant cycle of the GluN1-S688 residue. (**A**) Summary of the relative surface expression of NMDARs containing the indicated mutations of the GluN1-S688 residue, measured using fluorescence microscopy; \**p* < 0.050, and \*\*\**p* < 0.001 vs WT; one-way ANOVA. Data points correspond to individual cells (*n* ≥ 82), and the red box plot represents mean ± SEM. (**B**) Normalized concentration-response curves for glycine (gly) measured from HEK293 cells expressing WT and mutant NMDARs were obtained by fitting the data using Equation (1) (see Methods); for the summary of fitting parameters, see **Table 4**. **(C)** Correlation of surface expression and EC_50_ values for glycine for WT and mutated YFP-hGluN1-1a-S688/hGluN2A receptors containing indicated mutant cycle variants were fitted by linear regression. **(D-I)** Molecular modeling and molecular dynamics simulations depict human GluN1 LBD binding site complexes with glycine. Each figure illustrates a cartoon representation of the complex, including superimposed glycine **(D)**, GluN1-S688A-glycine complex **(E)**, GluN1-S688Y-glycine complex **(F)**, GluN1-S688K-glycine complex **(G)**, GluN1-S688C-glycine complex **(H)**, and GluN1-S688P-glycine complex **(I)**. Amino acid residues crucial for glycine binding are differentiated by rendering dark blue in orange and light blue in salmon. Polar contacts are indicated by black dashed lines. The structures are based on the PDB ID: 5KCJ.

To understand at the structural level how the selected amino acid substitutions altered the interaction of the LBDs with their principal agonists, we performed molecular modeling (MM) in tandem with molecular dynamics (MD) on the human GluN1/GluN2A LBD structure (PDB ID: 5KCJ). In case two equipotent amino acid mutant rotamer states were found, those are displayed and discussed accordingly. To validate our *in silico* protocol, we first calculated glycine binding to the GluN1 subunit (Figure 6D) and L-glutamate binding to the GluN2A subunit (Figure 8D), respectively. The root mean square deviation (RMSD) of all heavy atoms between the X-ray structure (PDB ID: 5KCJ) and the docked agonists with superimposed GluN1 and GluN2 subunits resulted in 0.749 Å^2^ and 0.671 Å^2^ values, respectively, indicating in both cases excellent performance with high model reliability. All pivotal hydrogen bond interactions between agonists and amino acid residues were preserved across LBDs of GluN1 and GluN2A subunits compared to crystal structure (Figure 6D and 8D). Water molecules were also taken into calculations, but for the sake of clarity, they were omitted from the graphical representations. Furthermore, using UCSF Chimera software, we generated the GluN1/GluN2A LBD structures with the mutant cycles of the GluN1-688, GluN1-D732, GluN2A-S511, and GluN2A-T690 residues, the resulting binding energies for glycine or L-glutamate obtained by MD simulations are shown in Table 5. From these data, it can be postulated that the correlation between EC_50_ values and binding energies from *in silico* is inconsistent and depends on several factors. First, it is essential to differentiate between EC_50_ and binding affinity. Fundamentally, while binding affinity is a major factor influencing EC_50_, it is not the sole determinant. The EC_50_ also depends on the downstream effects following the binding event, including cellular and systemic responses, which are beyond the scope of *in silico* studies focused solely on binding affinity. Thus, discrepancies between *in silico* binding affinity predictions and experimentally measured EC_50_ values can arise due to these additional complexities.

**Figure 8.**
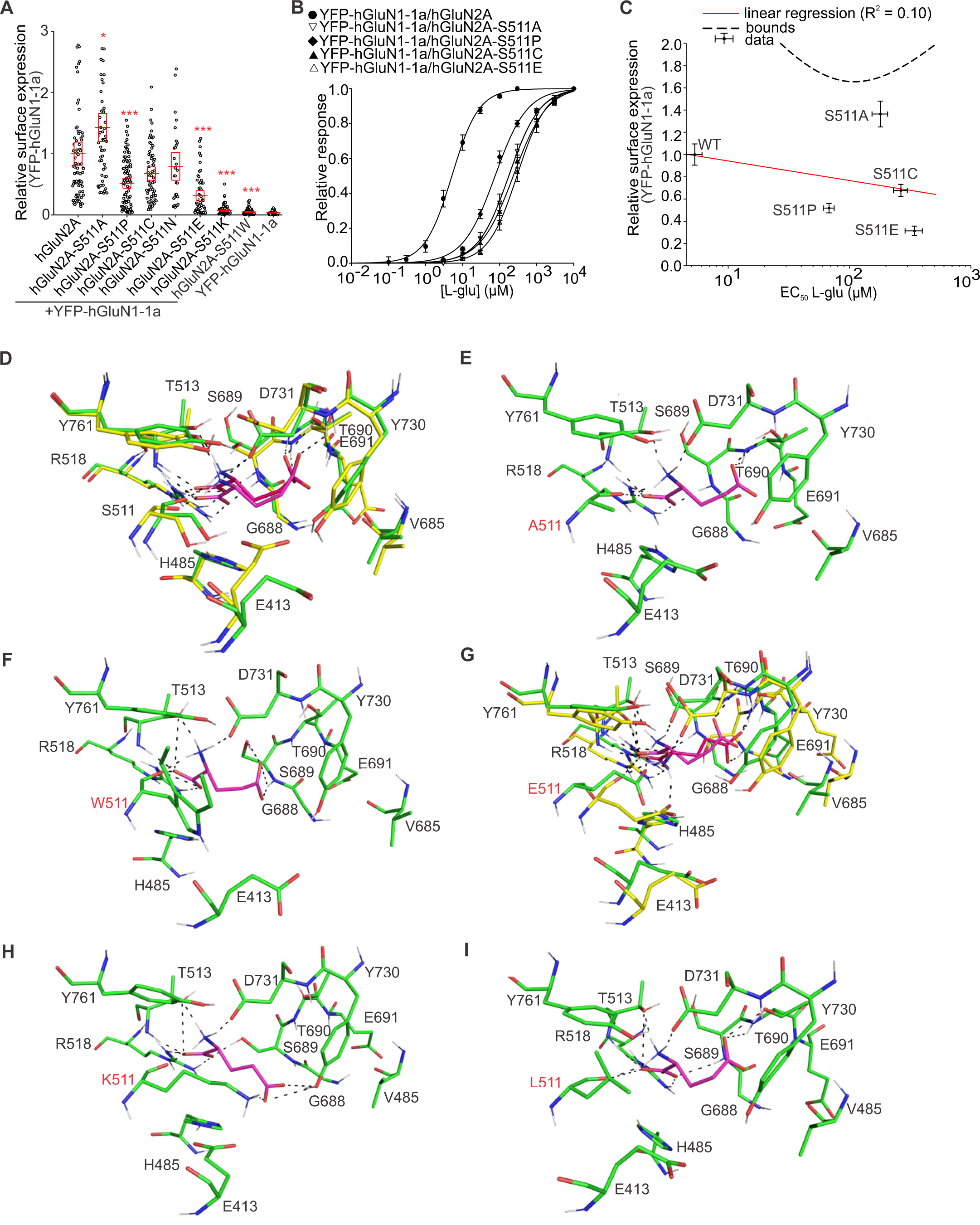
The mutant cycle of the GluN2A-S511 residue. **(A)** Summary of the relative surface expression of NMDARs containing the indicated mutations, measured using fluorescence microscopy; \**p* < 0.050, and \*\*\**p* < 0.001 vs WT; one-way ANOVA. Data points correspond to individual cells (*n* ≥ 29), and the red box plot represents mean ± SEM. (**B**) Normalized concentration-response curves for L-glutamate (L-glu) measured from HEK293 cells expressing NMDARs containing mutations of the GluN2A-S511 residue obtained by fitting the data using Equation (1) (see Methods); for the summary of fitting parameters, see **Table 4**. **(C)** Correlation of surface expression and EC_50_ values for glycine for WT and mutated YFP-hGluN1-1a/hGluN2A-S511 receptors containing indicated mutations fitted by linear regression. **(D-I)** Molecular modeling and molecular dynamics simulations depict human GluN2A LBD binding site complexes with L-glutamate. Each figure illustrates a cartoon representation of the complex, including superimposed L-glutamate **(D)**, GluN2A-S511A-L-glutamate complex **(E)**, GluN2A-S511W-L-glutamate complex **(F)**, GluN2A-S511E-L-glutamate complex **(G)**, GluN2A-S511K-L-glutamate complex (**H**), and GluN2A-S511L-L-glutamate complex **(I)**. Amino acid residues crucial for L-glutamate binding are differentiated by rendering green in purple and yellow in red. Polar contacts are indicated by black dashed lines. The structures are based on the PDB ID: 5KCJ.

**Table 5.**
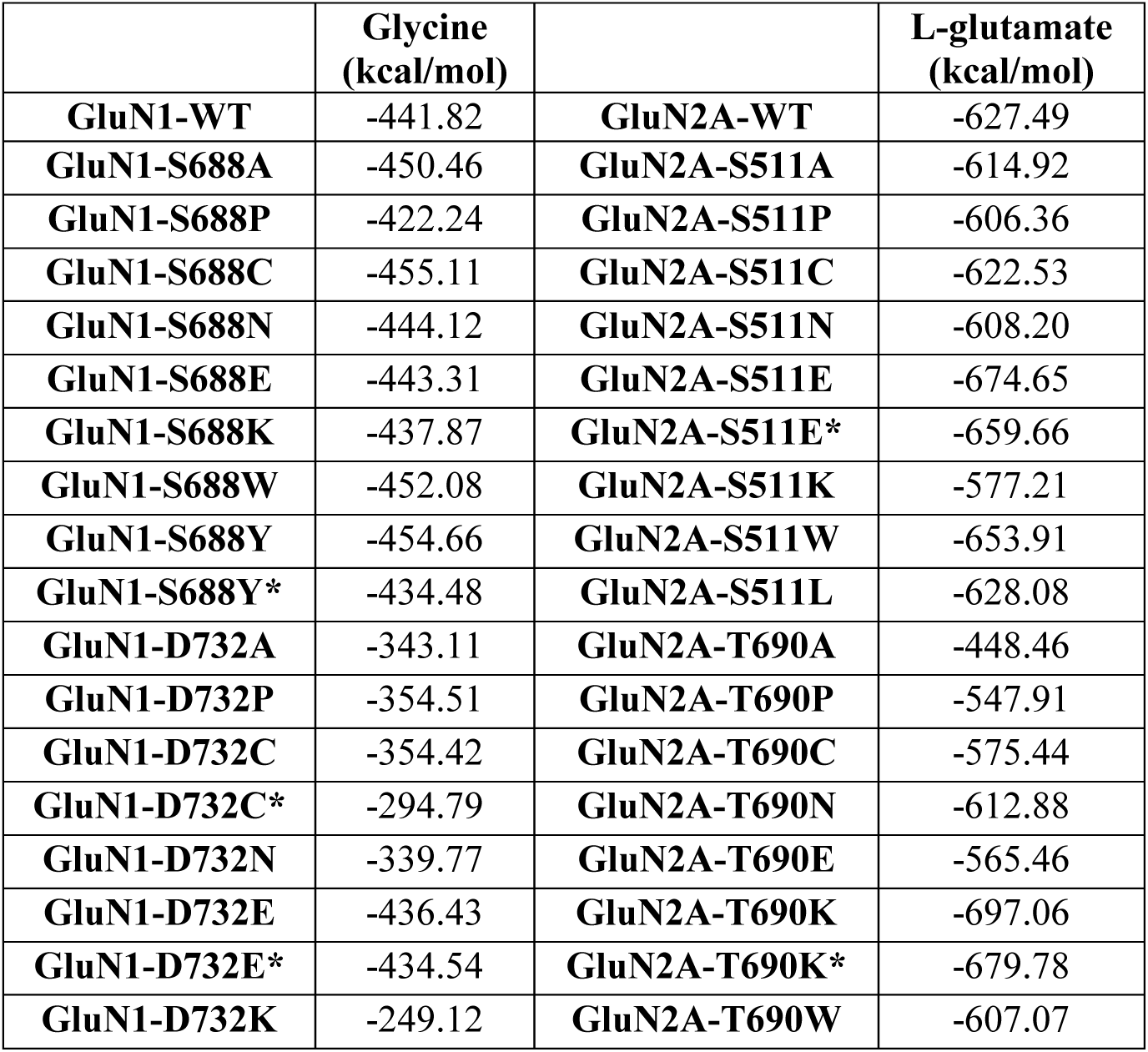
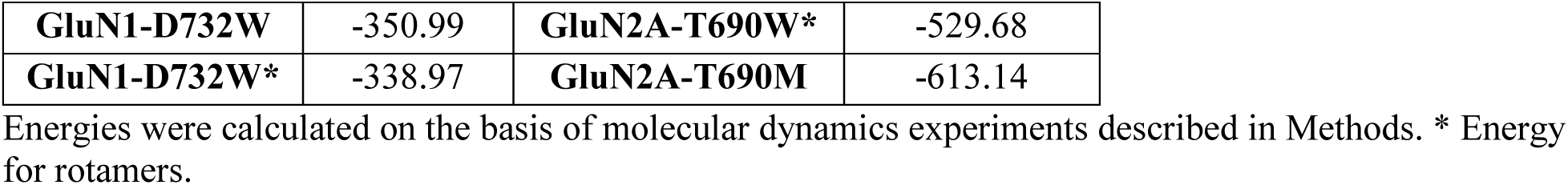
Binding energy to LBDs of GluN1 subunit for glycine and GluN2A subunit to L-glutamate obtained by molecular dynamic simulations.

|  | **Glycine**<br>**(kcal/mol)** |  | **L-glutamate**<br>**(kcal/mol)** |
| --- | --- | --- | --- |
| **GluN1-WT** | -441.82 | **GluN2A-WT** | -627.49 |
| **GluN1-S688A** | -450.46 | **GluN2A-S511A** | -614.92 |
| **GluN1-S688P** | -422.24 | **GluN2A-S511P** | -606.36 |
| **GluN1-S688C** | -455.11 | **GluN2A-S511C** | -622.53 |
| **GluN1-S688N** | -444.12 | **GluN2A-S511N** | -608.20 |
| **GluN1-S688E** | -443.31 | **GluN2A-S511E** | -674.65 |
| **GluN1-S688K** | -437.87 | **GluN2A-S511E\*** | -659.66 |
| **GluN1-S688W** | -452.08 | **GluN2A-S511K** | -577.21 |
| **GluN1-S688Y** | -454.66 | **GluN2A-S511W** | -653.91 |
| **GluN1-S688Y\*** | -434.48 | **GluN2A-S511L** | -628.08 |
| **GluN1-D732A** | -343.11 | **GluN2A-T690A** | -448.46 |
| **GluN1-D732P** | -354.51 | **GluN2A-T690P** | -547.91 |
| **GluN1-D732C** | -354.42 | **GluN2A-T690C** | -575.44 |
| **GluN1-D732C\*** | -294.79 | **GluN2A-T690N** | -612.88 |
| **GluN1-D732N** | -339.77 | **GluN2A-T690E** | -565.46 |
| **GluN1-D732E** | -436.43 | **GluN2A-T690K** | -697.06 |
| **GluN1-D732E\*** | -434.54 | **GluN2A-T690K\*** | -679.78 |
| **GluN1-D732K** | -249.12 | **GluN2A-T690W** | -607.07 |
| **GluN1-D732W** | -350.99 | **GluN2A-T690W\*** | -529.68 |
| **GluN1-D732W\*** | -338.97 | **GluN2A-T690M** | -613.14 |

Energies were calculated on the basis of molecular dynamics experiments described in Methods. \* Energy for rotamers.

Initially, we embarked on a detailed analysis of the mutant cycle of GluN1-S688 residue, specifically GluN1-S688A, GluN1-S688Y, GluN1-S688K, GluN1-S688C, and GluN1-S688P mutations. The rationale for the mutant selection reflects the prototypical alanine substitution (GluN1-S688A; Figure 6E), the highest surface expression and known pathogenicity (GluN1-S688Y; Figure 6F), the lowest surface expression (GluN1-S688K; Figure 6G), the highest (GluN1-S688C; Figure 6H, Table 5), and the lowest MM/MD energy scores and known pathogenicity (GluN1-S688P; Figure 6I, Table 5). The more negative docking scores for the MM/MD score indicate a higher agonist (glycine) affinity for the LBD of the GluN1 subunit. In all studied complexes, the basic hydrogen bonds were preserved: i) namely hydrogen bond tripod between glycine ammonium moiety with P516 amide, D732 carboxylate, and T518 hydroxyl, ii) cation-π interaction between glycine ammonium moiety and F484, and iii) salt bridge contacts between glycine carboxylate to GluN1-R523 guanidinium moiety, and NH amide backbone, except for S688P mutation. In the case of GluN1-S688K, GluN1-S688P, and GluN1-S688Y, more sterically demanding amino acid residues may affect the agonist binding, pushing the glycine out of the GluN1-R523 guanidinium moiety. Intriguingly, two possible GluN1-Y688 rotamer orientations are equally probable (Figure 6F), fixing the agonist similarly. Another study also observed the latter dealing with MM/MD simulation between glycine and the GluN1-S688Y subunit (Chen et al., 2023).

Regarding the mutant cycle of the GluN1-D732 residue, our microscopical analysis showed the following order for the surface expression levels of YFP-hGluN1-1a/hGluN2A receptors in the HEK293 cells: GluN1-D732W> WT> GluN1-D732E> GluN1-D732K> GluN1-D732P> GluN1-D732C> GluN1-D732N> GluN1-D732A (Figure 7A). Using electrophysiology, we did not detect any current responses of YFP-hGluN1-1a/hGluN2A receptors containing the GluN1-D732P and GluN1-D732W mutations, and we obtained current responses <100 pA for the YFP-hGluN1-1a-D732A/hGluN2A receptors (which could not be analyzed using logistic *Equation 1*; Figure 7B). For the other mutant YFP-hGluN1-1a/hGluN2A receptors, we calculated their EC_50_ values for glycine, which very weakly negatively correlated (R^2^ = 0.10) with the surface expression levels (Figure 7C). Furthermore, we employed MM with MD, focusing on the GluN1-D732W displaying the highest surface expression (Figure 7D), and the GluN1-D732A mutation demonstrated the lowest surface expression and prototypical alanine substitution (Figure 7E). In addition, we examined both GluN1-D732E (Figure 7F) and GluN1-D732K (Figure 7G) pathogenic variants, which also exhibited the highest and the lowest MM/MD energy scores, respectively (Table 5). The two most populated rotamer clusters were observed for GluN1-D732E and GluN1-D732W mutations, giving rise to the equipotent conformational probability of the complex. For the GluN1-D732E variant, both complexes were almost identical (RMSD = 1.034 Å^2^) with overlapping GluN1-E732 residue, while GluN1-W732 residue revealed two distinct orientations (RMSD = 1.017 Å^2^). For GluN1-D732E, GluN1-D732W, and GluN1-D732A complexes, interactions between carboxylate glycine and GluN1-R532 guanidinium moiety, GluN1-S688 NH amide backbone, and GluN1-T518 hydroxyl were well-conserved. The positively charged ammonium moieties between glycine and GluN1-K732 in GluN1-D732K structure shifted the agonist to a less favorable position by repulse interaction, with glycine ammonium moiety facing GluN1-T518 hydroxyl, concomitantly weakening cation-π stacking with GluN1-F484 residue. Notably, the salt bridge interaction between glycine ammonium and GluN1-W732 and GluN1-A732 residues was missing.

**Figure 7.**
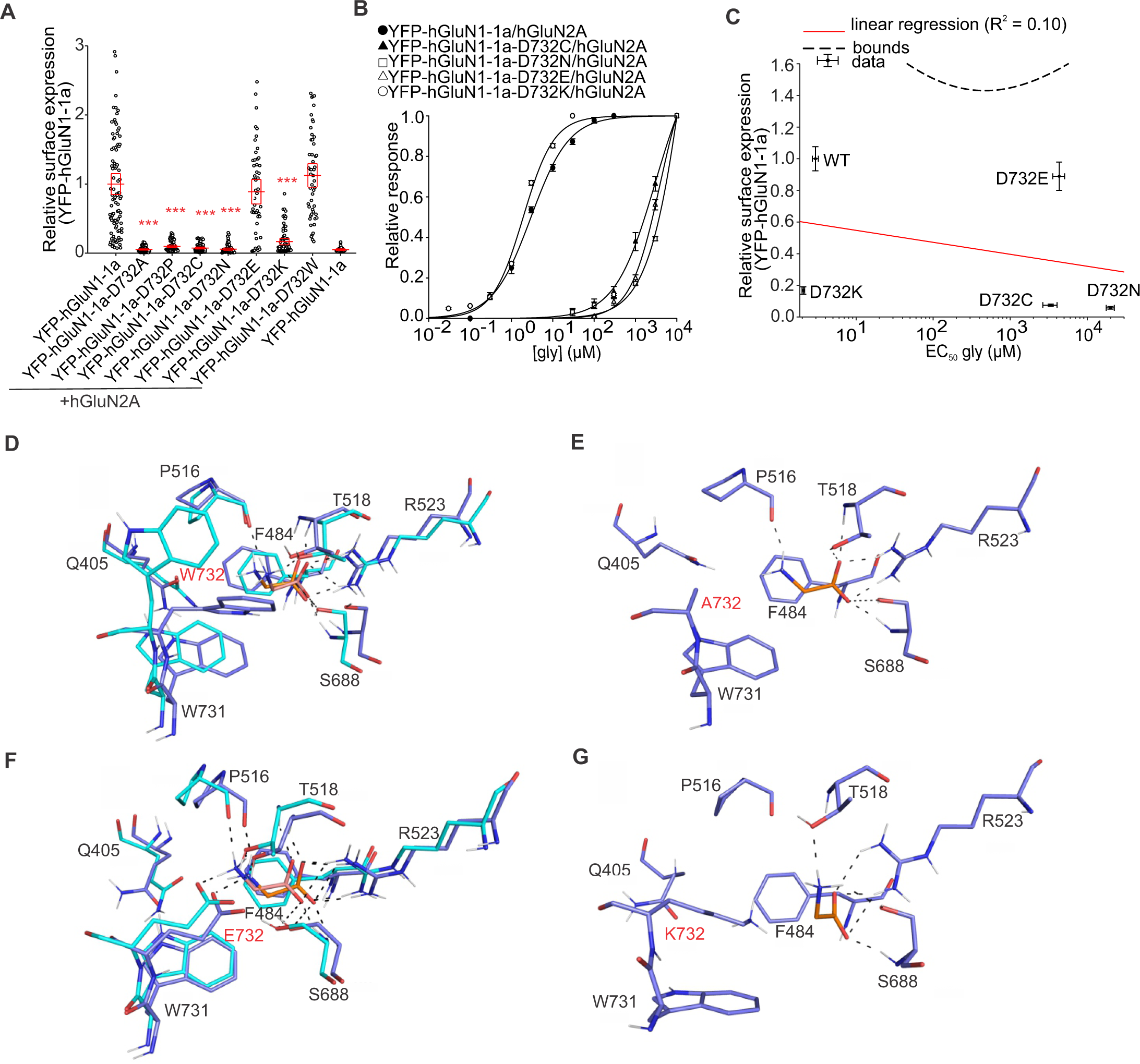
The mutant cycle of the hGluN1-D732 residue. **(A)** Summary of the relative surface expression of NMDARs containing the indicated mutations, measured using fluorescence microscopy; \*\*\**p* < 0.001 vs WT; one-way ANOVA. Data points correspond to individual cells (*n* ≥ 34), and the red box plot represents mean ± SEM. **(B)** Normalized concentration-response curves for glycine (gly) measured from HEK293 cells expressing NMDARs containing mutations of the GluN1-D732 residue obtained by fitting the data using Equation (1) (see Methods); for the summary of fitting parameters, see **Table 4**. **(C)** Correlation of surface expression and EC_50_ values for glycine for WT and mutated YFP-hGluN1-1a-D732/hGluN2A receptors containing indicated mutations fitted by linear regression. **(D-G)** Molecular modeling and molecular dynamics simulations depict human GluN1 LBD binding site complexes with glycine. Each figure illustrates a cartoon representation of the complex, including GluN1-D732W-glycine complex **(D)**, GluN1-D732A-glycine complex **(E)**, GluN1-D732E-glycine complex **(F)**, and GluN1-D732K-glycine complex **(G)**. Amino acid residues crucial for glycine binding are differentiated by rendering dark blue in orange and light blue in salmon. Polar contacts are indicated by black dashed lines. The structures are based on the PDB ID: 5KCJ.

Regarding the mutation cycle of the GluN2A-S511 residue, we observed the following trend in the surface expression of YFP-hGluN1-1a/hGluN2A receptors in HEK293 cells: GluN2A-S511A>WT> GluN2A-S511N> GluN2A-S511C> GluN2A-S511P> GluN2A-S511E> GluN2A-S511K> GluN2A-S511W (Figure 8A). Our electrophysiological measurements from HEK293 cells expressing YFP-hGluN1-1a/hGluN2A receptors containing GluN2A-S511N, GluN2A-S511K, GluN2A-S511W mutations showed no current responses; therefore, we calculated EC_50_ values for L-glutamate for the remaining mutant YFP-hGluN1-1a/hGluN2A receptors (Figure 8B). Using linear regression, we observed a very weak negative correlation (R^2^ = 0.10) between the surface expression levels and the EC_50_ values for L-glutamate of the YFP-hGluN1-1a/hGluN2A receptors with the mutant cycle at the GluN2A-T690 position (Figure 8C). We also gained structural insight into L-glutamate binding to GluN2A-S511A (Figure 8E; the highest surface expression), GluN2A-S511W (Figure 8F; the lowest surface expression), GluN2A-S511E (Figure 8G; the highest MM/MD energy score, Table 5), GluN2A-S511K (Figure 8H; the lowest MM/MD energy score, Table 5), and GluN2A-S511L (Figure 8I; pathogenic variant). For GluN2A-S511A, ammonium glycine moiety lacks contact with GluN2A-A511 residue, while all other salt bridge interactions are conserved compared to crystal L-glutamate structure. Interestingly, in the L-glutamate/GluN2A-S511W complex, all hydrogen contacts were also preserved with tight conformation between the carbonyl amide of GluN2A-W511 residue and ammonium L-glutamate moiety, further displacing the γ-carboxylate moiety outwards from amide backbone GluN2A-T690 and GluN2A-E691 residues, the interactions typically occurring in the WT L-glutamate/GluN2A anchoring. In the GluN2A-S511E mutation, the two highest equally possible rotamer clusters were possible, which are outlined accordingly. Interestingly, both GluN2A-E511 rotamers can attract L-glutamate ammonium moiety to carboxylate residue. In the case of L-glutamate depicted in purple, these attractive forces resulted in the loss of the salt bridge to GluN2A-S689 residue. Completely different hydrogen bond network distribution can be observed for the GluN2A-S511K mutation, as the side chain of GluN2A-K511 residue deflected γ-carboxylate L-glutamate from its native conformation, completely losing contact with GluN2A-S689 and GluN2A-T690 residues. Under these circumstances, polar contact to Y730 hydroxyl moiety has been formed. For the WT LBD of the GluN2A subunit, GluN2A-Y730 residue is one of the amino acid residues directly implicated in contact with L-glutamate via hydrophobic interactions. No specific changes compared to the WT L-glutamate/GluN2A complex were found for the GluN2A-S511L residue.

Finally, we observed the following trend in the surface expression of YFP-hGluN1-1a/hGluN2A receptors with the mutant cycle of the GluN2A-T690 residue: WT> GluN2A-T690C> GluN2A-T690A> GluN2A-T690N> GluN2A-T690P> GluN2A-T690E> GluN2A-T690W> GluN2A-T690K (Figure 9A). We observed no current responses using electrophysiology for YFP-hGluN1-1a/hGluN2A receptors containing the GluN2A-T690P, GluN2A-T690N, GluN2A-T690E, GluN2A-T690K and GluN2A-T690W subunits, nor with the mutant hGluN1-4a subunit (data not shown; Figure 9B). Therefore, we further calculated EC_50_ values for L-glutamate only for YFP-hGluN1-1a/hGluN2A receptors containing the GluN2A-T690C and GluN2A-T690A mutation (Figure 4) and compared them by linear regression with surface expression levels; which showed a very strong negative correlation (R^2^ = 1.00, Figure 9C). We also performed MM/MD experiments on the mutant cycle of the GluN2A-T690 residue. Specifically, we studied GluN2A-T690K (highest MM/MD energy score; Figure 9D), GluN2A-T690A (Figure 9E; lowest MM/MD energy score, Table 5), GluN2A-T690C (Figure 9F; highest surface expression), GluN2A-T690W (Figure 9G; lowest surface expression) and GluN2A-T690M (Figure 9H; pathogenic variant) mutations. Two rotamer conformations were possible in the case of GluN2A-T690K and GluN2A-T690W mutations. For the L-glutamate/T690K complex, no structural deviations from WT GluN2A subunit were found; L-glutamate γ-carboxylate revealed typical hydrogen bonds with amide backbones of GluN2A-S689 and GluN2A-K690 residues. However, the presence of GluN2A-K690 residue likely affected the distance between the L-glutamate and GluN2A-Y730 residue employed in the hydrophobic contact with the L-glutamate. Hydrogen bond contact between L-glutamate and amide backbone from GluN2A-D731 residue was also absent. In the L-glutamate/GluN2A-T690A complex, the GluN2A-A690 residue caused slightly different L-glutamate accommodation than the native state. Indeed, the γ-carboxylate protrudes out of GluN2A-D731 and GluN2A-Y730 residues, enabling the move of GluN2A-D731 carboxylate into the proximity of L-glutamate ammonium moiety. This structural conformation seems well-corroborated by salt bridge interactions with the amide backbones of GluN2A-A690 and GluN2A-E691 residues. Interestingly, the L-glutamate/GluN2A-T690C complex yielded a more distanced topology of γ-carboxylate from GluN2A-C690 residues, fixing the moiety via a double-mediated hydrogen bond to S689 amide and 689 hydroxyl. In other words, the GluN2A-C690 residue was not involved in anchoring L-glutamate directly. Two completely different clusters were gained for the GluN2A-T690W mutation. The rotamer cluster with GluN2A-W690 depicted in purple (Figure 9G) generated U-shaped L-glutamate binding centered around GluN2A-S689 residue. Thus, GluN2A-W690 residue impacted the L-glutamate binding in less energetically favorable conformation via its sterical effect and negatively charged area, given its electron-rich character. More strikingly, the second rotamer conformation (highlighted with red GluN2A-W690 residue, Figure 9G) had the same negative effect, affecting L-glutamate spatial orientation into linear form, with a γ-carboxylate orientation towards the hydroxyl moiety of GluN2A-Y730 residue, and with no GluN2A-E691 residue contact. In the L-glutamate/GluN2A-T690M complex, no deviations compared to crystal L-glutamate crystal structure were apparent except for closer γ-carboxylate binding to amide backbones of GluN2A-S689 and GluN2A-M690 residues along with GluN2A-S689 hydroxyl moiety hydrogen bond on the cost of more distanced GluN2A-Y730 hydrophobic contact.

**Figure 9.**
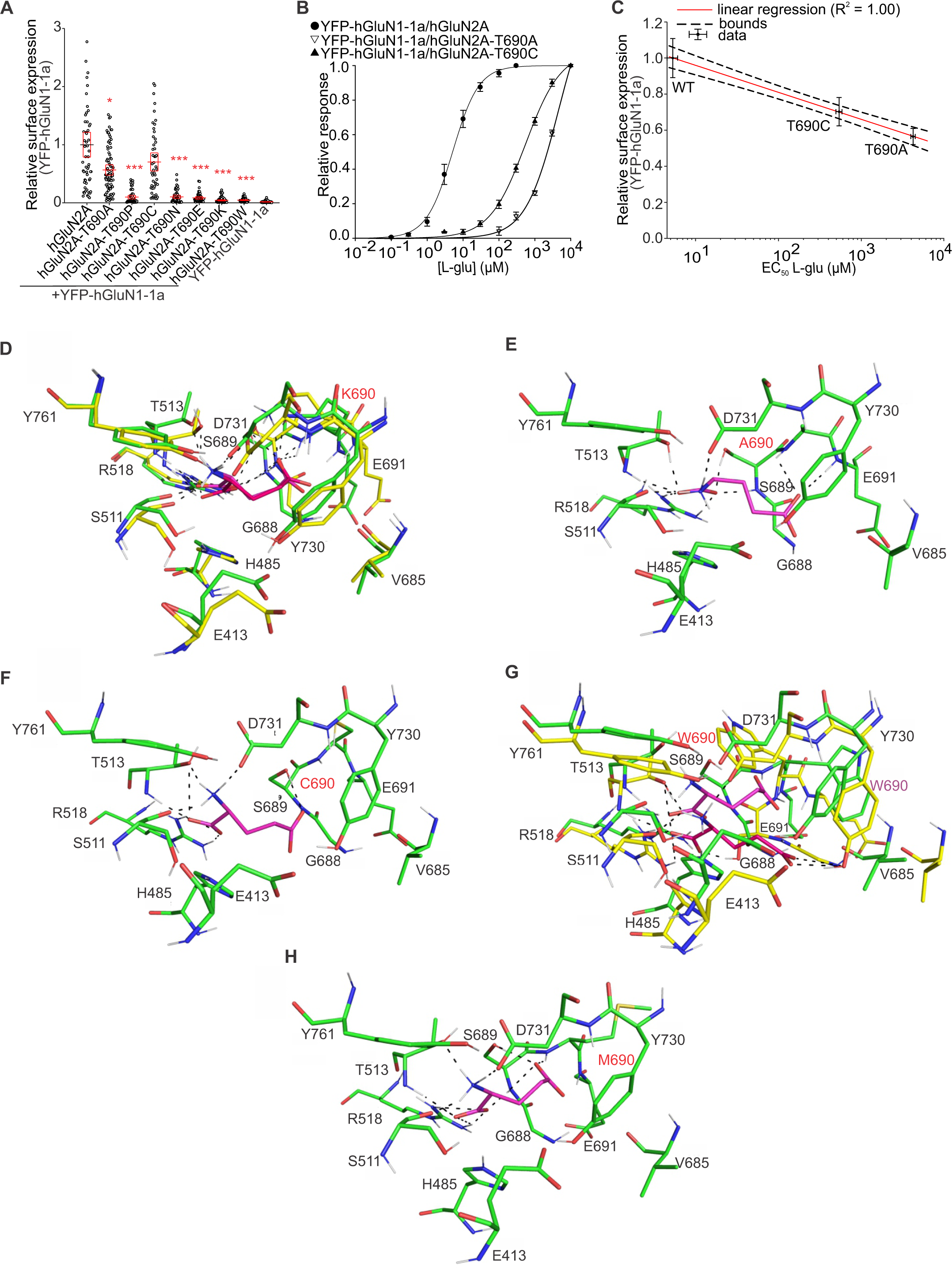
The mutant cycle of the GluN2A-T690 residue. **(A)** Summary of the relative surface expression of NMDARs containing the indicated mutations of the GluN2A-T690 residue, measured using fluorescence microscopy; \**p* < 0.050, and \*\*\**p* < 0.001 vs WT; one-way ANOVA. Data points correspond to individual cells (*n* ≥ 49), and the red box plot represents mean ± SEM. (**B**) Normalized concentration-response curves for L-glutamate (L-glu) measured from HEK293 cells expressing mutant NMDARs obtained by fitting the data using Equation (1) (see Methods); for the summary of fitting parameters, see **Table 4**. **(C)** Correlation of surface expression and EC_50_ values for glycine for WT and mutated YFP-hGluN1-1a/hGluN2A receptors fitted by linear regression. **(D-H)** Molecular modeling and molecular dynamics simulations depict human GluN2A LBD binding site complexes with L-glutamate. Each figure illustrates a cartoon representation of the complex, including GluN2A-T690K-L-glutamate complex **(D)**, GluN2A-T690A-L-glutamate complex **(E)**, GluN2A-T690C-L-glutamate complex **(F)**, GluN2A-T690W-L-glutamate complex with highlighted rotamers in red and purple **(G)**, and GluN2A-T690M-L-glutamate complex **(H)**. Amino acid residues crucial for L-glutamate binding are differentiated by rendering green in purple and yellow in red. Black dashed lines indicate polar contacts. The structures are based on the PDB ID: 5KCJ.

### The effect of selected alanine substitutions in LBDs on surface expression is conserved for GluN1/GluN2A and GluN1/GluN2B receptors

We next asked whether the selected alanine substitutions in LBD of the GluN1 subunit, GluN1-F484A, GluN1-P516A, GluN1-R523A GluN1-S688A and GluN1-D732A, examined in more detail in combination with the GluN2A subunit, alter similarly the surface expression of the GluN1/GluN2B receptors. Using immunostaining with an anti-GFP antibody, we observed that the relative surface expression levels of the YFP-GluN1-1a/GluN2B receptors showed a similar order to that of the YFP-GluN1-1a/GluN2A receptors, specifically WT > GluN1-S688A > GluN1-F484A > GluN1-P516A > GluN1-D732A > GluN1-R523A (Figure 10A). After plotting the surface expression values with linear regression, we obtained a very strong positive correlation (R^2^ = 0.91) between mutant YFP-GluN1-1a/GluN2A and YFP-GluN1-1a/GluN2B receptors, indicating that the surface expression of both subtypes is similarly regulated by structural changes in LBD of the GluN1 subunit (Figure 10B).

**Figure 10.**
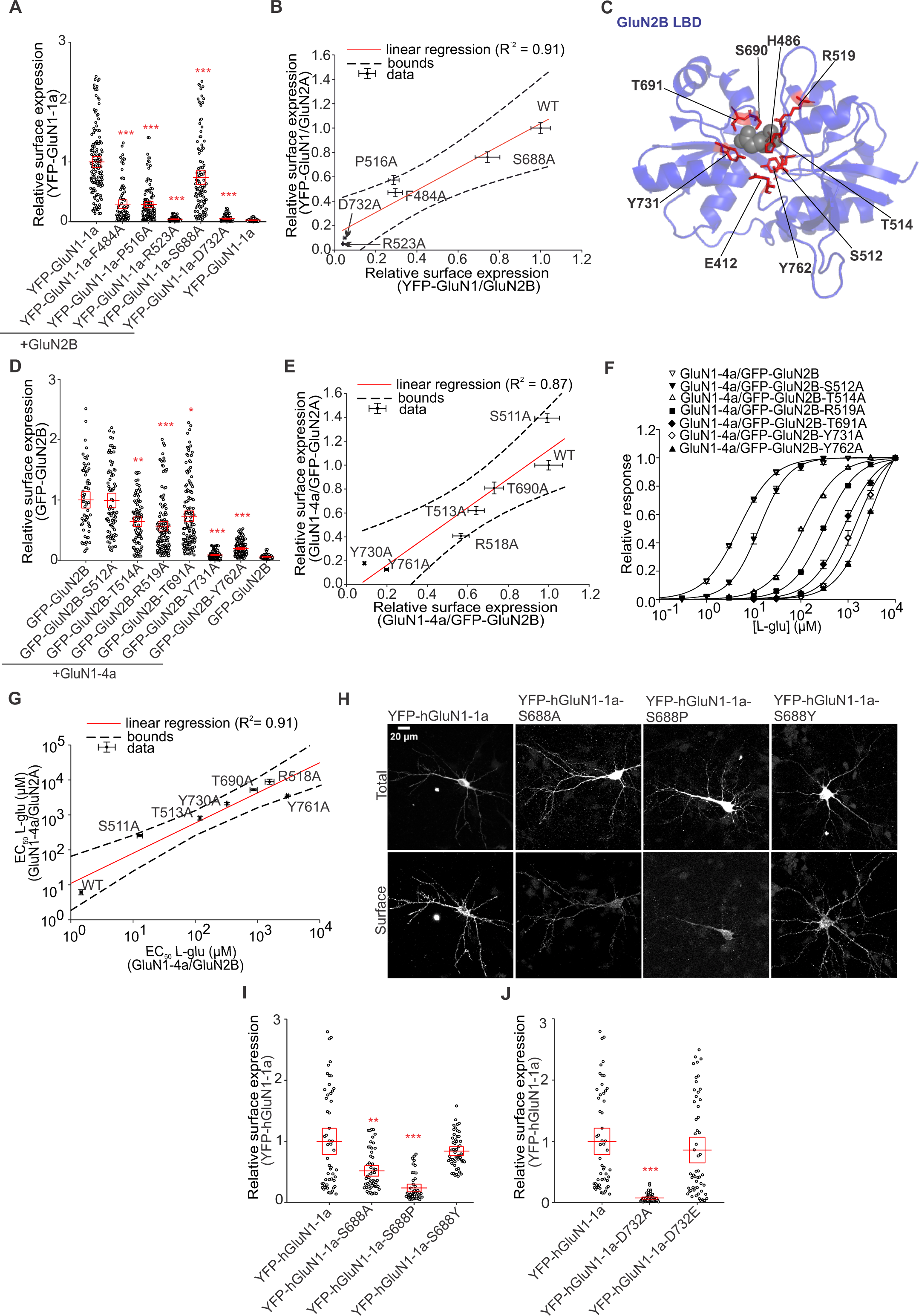
Alanine substitutions in LBDs cause similar changes in surface expression of GluN1/GluN2B receptors as we observed for GluN1/GluN2A receptors. **(A)** Summary of the relative surface expression of NMDARs consisting of either WT or mutated YFP-GluN1-1a subunit co-expressed with the GluN2B subunit, measured using fluorescence microscopy; ***p < 0.001 vs WT, one-way ANOVA. Data points correspond to individual cells (*n* ≥ 84), and the red box plot represents mean ± SEM. **(B)** Correlation of surface expression of YFP-GluN1-1a/GluN2A and YFP-GluN1-1a/GluN2B receptors. **(C)** Structural model of the LBD of the GluN2B subunit labeled with L-glutamate-interacting amino acids (based on PDB ID: 4PE5). **(D)** Summary of the relative surface expression of NMDARs consisting of either WT or mutated GFP-GluN2B subunit co-expressed with the GluN1-4a subunit, measured using fluorescence microscopy; *p < 0.050, **p < 0.010, and ***p < 0.001 vs WT; one-way ANOVA. Data points correspond to individual cells (*n* ≥ 64), and the red box plot represents mean ± SEM. **(E)** Correlation between surface expression levels of homologous mutations in GluN2A and GluN2B subunits when co-expressed with GluN1-4a subunit. The plotted points in the graph were labeled according to the amino acid positions of the GluN2A subunit. **(F)** Normalized concentration-response curves for L-glutamate effect on NMDARs containing the indicated mutation obtained by fitting the data using Equation (1) (see Methods); for a summary of fitting parameters, see Table 6. **(G)** Correlation between EC_50_ values measured for WT and mutated GluN1/GluN2A and GluN1/GluN2B receptors. The plotted points in the graph were labeled according to the amino acid positions of the GluN2A subunit. **(H)** Representative images of hippocampal neurons transfected with the WT or mutated YFP-hGluN1-1a subunits. The total and the surface signal (top and bottom row, respectively) of YFP-GluN1-1a subunits were labeled using an anti-GFP antibody 48 hours after the transfection. **(I,J)** Summary of the relative surface expression of YFP-hGluN1-1a subunits mutated at position S688 **(I)** or D732 (**J**), expressed in hippocampal neurons determined using fluorescence microscopy; \*\*\**p* < 0.001, \*\**p* < 0.010 vs WT; one-way ANOVA. Data points correspond to individual segments (*n* ≥ 46), and the red box plot represents mean ± SEM.

**Table 6.** Summary of fitting parameters for steady-state concentration-response curves for L-glutamate measured in HEK293 cells expressing the indicated GluN1/GluN2B receptors.

| **Receptor** | **EC<sub>50</sub> (μM)** | ***h*** | ***n*** |
| --- | --- | --- | --- |
| **GluN1-4a/**<br>**GFP-GluN2B** | 1.46 ± 0.07 | 1.22 ± 0.07 | 9 |
| **GluN1-4a/**<br>**GFP-GluN2B-S512A** | 12.80 ± 1.57 | 1.43 ± 0.06 | 6 |
| **GluN1-4a/**<br>**GFP-GluN2B-T514A** | 118.58 ± 4.97 | 1.14 ± 0.03 | 7 |
| **GluN1-4a/**<br>**GFP-GluN2B-R519A** | 1,562.84 ± 263.28 | 1.35 ± 0.07 | 4 |
| **GluN1-4a/**<br>**GFP-GluN2B-T691A** | 867.10 ± 124.90 | 1.24 ± 0.06 | 5 |
| **GluN1-4a/**<br>**GFP-GluN2B-Y731A** | 324.94 ± 9.68 | 1.19 ± 0.03 | 9 |
| **GluN1-4a/**<br>**GFP-GluN2B-Y762A** | 3,053.74 ± 155.64 | 1.22 ± 0.05 | 5 |

The EC<sub>50</sub> and *h* values were obtained by fitting the data from individual cells to Equation (1).

We further investigated whether the surface expression of GluN1/GluN2B receptors is regulated by structural changes in the LBD of the GluN2B subunit, similar to the case of GluN1/GluN2A receptors. The structure of the LBD of the GluN2B subunit with labeled amino acid residues directly interacting with L-glutamate is shown in Figure 10C; interestingly, the amino acid residues interacting with L-glutamate are conserved in all GluN2 subunits (Kinarsky et al., 2005). Therefore, we further compared the surface expression of the GluN1-4a/GFP-GluN2B receptors containing the corresponding alanine substitutions, GluN2B-S512A, GluN2B-T514A, GluN2B-R519A, GluN2B-T691A, GluN2B-Y731A, and GluN2B-Y761A, which we studied in detail in the case of GluN1/GluN2A receptors. Analysis of the surface fluorescence signals of the GluN1-4a/GFP-GluN2B receptors expressed in the HEK293 cells showed the following order: WT> GluN2B-S512A> GluN2B-T691A> GluN2B-T514A> GluN2B-R519A> GluN2B-Y762A> GluN2B-Y731A (Figure 10D); which, except for mutations causing highly deficient surface expression (GluN2B-R519A, GluN2B-Y761A, GluN2B-Y731A), corresponded to the order obtained for homologous mutations in the case of the GluN2A subunit (see Figure 2). This conclusion was confirmed by the very strong positive correlation (R^2^ = 0.87) between the surface expression of the GluN1/GluN2A and GluN1/GluN2B receptors containing the corresponding alanine substitutions within the LBDs of the GluN2A and GluN2B subunits, respectively (Figure 10E). Our subsequent electrophysiological recordings and concentration-dependent analysis of WT and mutant GluN1-4a/GFP-GluN2B receptors revealed a very strongly positive correlation (R^2^ = 0.91) between the EC_50_ values for L-glutamate measured for the GluN1/GluN2A and GluN1/GluN2B receptors (Figure 10F-G; Table 6), showing that homologous alanine substitutions in LBDs of the GluN2A and GluN2B subunits have similar effects on the sensitivity of the mutated GluN1/GluN2 receptors to L-glutamate.

Finally, we asked whether our findings on the role of LBDs in regulating the surface number of NMDARs gained using HEK293 cells also apply to hippocampal neurons. Therefore, we determined the surface expression of YFP-hGluN1-1a subunits carrying alanine substitutions and pathogenic variants by transfecting them into cultured hippocampal neurons (DIV12) and labeling them using an anti-GFP antibody after 48 hours (Figure 10H). Our microscopic analysis showed the following decrease in surface expression of mutant YFP-hGluN1-1a subunits compared to the WT YFP-hGluN1-1a subunit: YFP-hGluN1-S688A (∼48% compared to ∼23% in HEK293 cells), YFP-hGluN1-S688P (∼76% compared to ∼59% in HEK293 cells), YFP-hGluN1-1a-D732A (∼93% compared to ∼94% in HEK293 cells). Similar to HEK293 cells, the YFP-hGluN1-S688Y and YFP-hGluN1-1a-D732E subunits expressed in hippocampal neurons did not exhibit altered surface expression compared to the WT YFP-GluN1-1a subunit. We could not verify the mutant GluN2A subunits in hippocampal neurons because we do not have a functional tagged human variant of the GluN2A subunit. These findings support the relevance of our conclusions obtained using HEK293 cells.

## Discussion

Previous studies of early trafficking of GluN1/GluN2 receptors studied a limited number of alanine substitutions that directly interact with glycine in the GluN1 subunit or L-glutamate in the GluN2A subunit. Specifically, the GluN1-D732A/GluN2A receptor showed a ∼90% reduction in surface expression, which was associated with a ∼35,000-fold increase in the EC_50_ value for glycine measured for the GluN1-D732A/GluN2B receptor expressed in *Xenopus* oocytes (Williams et al., 1996; Kenny et al., 2009). Our experiments with the GluN1-D732A/GluN2A receptor also showed a ∼90% reduction in surface expression, although we did not detect any current responses from transfected HEK293 cells. Previously, the GluN1/GluN2B-E413A receptor showed a ∼80% reduction in surface expression (She et al., 2012) and a ∼237-fold increase in EC_50_ value for L-glutamate (Laube et al., 2004). Consistent with this, our data with the GluN1/GluN2A-E413A receptor showed ∼80% reduction in surface expression and ∼287-fold increase in EC_50_ value for L-glutamate. In addition, EC_50_ values were also previously calculated for the following mutants of GluN1/GluN2A receptors: GluN1-F484A/GluN2A (glycine: 3,100 μM versus our measured value of 3,624 μM) (Kvist et al., 2013), GluN1-R523A/GluN2A (glycine: 5,400 μM versus our measured value of 10,261 μM) (Kvist et al., 2013), GluN1/GluN2A-S511A (L-glutamate: 146 μM versus our measurement of 235 μM) (Chen et al., 2005), GluN1/GluN2A-H485A (L-glutamate: 490-1,025 μM versus our measurement of 3,969 μM) (Anson et al., 1998; Chen et al., 2005; Maier et al., 2007), GluN1/GluN2A-T513A (L-glutamate: 482 μM versus our measurement of 809 μM) (Chen et al., 2005), GluN1/GluN2A-R518A (L-glutamate: >3000 μM versus our measurement of 8,895 μM) (Erreger et al., 2007), GluN1/GluN2A-S689A (L-glutamate: ∼2-5 μM versus our measurement of 7 μM) (Anson et al., 1998; Erreger et al., 2007) and GluN1/GluN2A-T690A (L-glutamate: 2,967 μM versus our measurement of 5,254 μM) (Anson et al., 1998). The observed differences in EC_50_ values can be explained by the use of different expression systems (*Xenopus* oocytes versus HEK293 cells), electrophysiological methods (two-electrode voltage clamp versus whole-cell patch-clamp), and differences in the composition of the recording solutions. Our series of alanine substitutions showed a strong (mutant GluN1 subunit) or weak (mutant GluN2A subunit) correlation between surface expression levels of GluN1/GluN2A receptors and their EC_50_ values for glycine or L-glutamate. Thus, our data showed that replacing critical amino acid residues within the LBDs of both the GluN1 and GluN2A subunits with a small uncharged alanine supports the previously hypothesized link between surface expression levels and the agonist potency of GluN1/GluN2 receptors. In addition, we found that co-expression of both GluN1 and GluN2A subunits with alanine substitutions in LBDs exhibited an additive effect on reducing the surface expression level of GluN1/GluN2A receptors, leading to the conclusion that LBDs of GluN1 and GluN2 subunits contribute independently to the regulation of the surface numbers of NMDARs. Our experiments with the closed cleft conformation of LBDs (Blanke and VanDongen, 2008; Kussius and Popescu, 2010; Dai and Zhou, 2016) confirmed the additive effect of the LBDs of the GluN1 and GluN2A subunits on the regulation of the surface numbers of the GluN1/GluN2A receptors. Our subsequent experiments revealed that closed cleft conformation of the LBDs of GluN1 and GluN2A subunits did not mask the reducing effect of alanine substitutions on the surface numbers of GluN1/GluN2A receptors. We further showed that incubating the transfected HEK293 cells with a panel of competitive antagonists did not alter the surface numbers of WT GluN1/GluN2A receptors. This result is consistent with previous data on the effect of DCKA on the early trafficking of WT GluN1/GluN2A receptor (Kenny et al., 2009). However, we cannot exclude that the selected competitive antagonists did not reach sufficient concentrations in the intracellular compartments, including the ER. Our experiments with alanine substitutions in the LBDs of GluN1/GluN2B receptors revealed similar findings to those observed with GluN1/GluN2A receptors, which suggests an evolutionarily shared mechanism for the regulation of the early trafficking of these two major diheteromeric subtypes of NMDARs.

Using the synchronized release of NMDARs from the ER, we showed that GluN1/GluN3A receptors co-localized with GA structures, similar to GluA1 receptors (Hangen et al., 2018). Consistently with a previous study showing diffuse localization of GluN1/GluN2A receptors in HEK293 cells (Hayashi et al., 2009), we did not detect GluN1/GluN2A receptors in GA, even at 15, 30, 45, and 60 min after addition of AL. Thus, the GluN1/GluN2A receptor may bypass the GA, as shown in a previous study with a transfected GluN1 subunit (Jeyifous et al., 2009). It was previously demonstrated that the WT GluN1/GluN2B receptor co-localizes with GA in COS-7 cells (She et al., 2012). Our pilot experiment did not detect WT GluN1/GluN2B receptors in GA structures using the ARIAD system; this discrepancy may be explained by the fact that expression of recombinant GluN1/GluN2B receptors may cause defragmentation of GA structures (Nakagomi et al., 2008) or cause excitotoxic damage (Zhou et al., 2013), which could affect the rate of co-localization between GA and NMDARs.

Regarding non-alanine substitutions in the GluN1 subunit, a previous study reported that the GluN1-D732E/GluN2A receptor did not show altered surface expression while exhibiting a ∼1,796-fold increase in EC_50_ value for glycine (Williams et al., 1996; Kenny et al., 2009), consistently with our results showing no difference in surface expression and ∼1,457-fold increase of EC_50_ value for glycin. In addition, our recent study reported that the pathogenic GluN1-S688Y variant did not alter the surface expression of the GluN1/GluN2A receptor but increased the EC_50_ values for glycine to 2,122 μM (Skrenkova et al., 2020), consistently with our novel data. In addition, our electrophysiological data with the GluN1-D732N/GluN2A receptor (EC_50_ values for glycine increased <4,000-fold compared with our measurement of 6,745-fold) (Williams et al., 1996; Seljeset et al., 2023) and the GluN1-D732W/GluN2A receptor (no current responses) (Seljeset et al., 2023) are consistent with previous studies. Regarding the published data with non-alanine substitutions in the GluN2A subunit, a previous study reported a 50% decrease, and we observed an 88% decrease in surface expression of the GluN1/GluN2A-R518H receptor (this difference may be explained by the use of colorimetric versus microscopic assays). Neither study detected current responses in HEK293 cells transfected with GluN1/GluN2A-R518H receptors (Swanger et al., 2016). In addition, the pathogenic GluN2B-E413G variant reduced surface expression of the GluN1/GluN2B receptor by ∼82% percent and increased the EC_50_ value for L-glutamate by ∼237-fold, which is consistent with our data with the GluN1/GluN2A-E413A receptor showing 71% percent lower surface expression and ∼288-fold higher EC_50_ value for L-glutamate. Our mutant cycles of GluN1-S688 and GluN1-D732 residues showed distinct levels of surface expression of GluN1/GluN2A receptors that did not correlate or correlated very weakly with EC_50_ values for glycine. In addition, mutant cycles at residues GluN2A-S511 and GluN2A-T690 showed strikingly different levels of surface expression, as they correlated weakly (GluN2A-S511 residue) or strongly (GluN2A-T690 residue, but only in the case of the two functional mutant GluN1/GluN2A receptors) with EC_50_ values for L-glutamate. Thus, non-alanine substitutions in LBDs caused changes in the surface expression of the GluN1/GluN2A receptor, which, unlike alanine substitutions, mostly did not correlate with EC_50_ values for glycine or L-glutamate. We hypothesized that the specific structural changes of LBDs, but not the functional properties, are the critical parameters sensed during the early trafficking of the GluN1/GluN2A receptor. This hypothesis is consistent with our observations that (i) the GluN1-D732E/GluN2A receptor was present on the cell surface similar to the WT GluN1/GluN2A receptor but had a ∼1,457-fold increased EC_50_ value for glycine, (ii) the GluN1/GluN2A-receptor showed a ∼43% increase of surface expression but a ∼34-fold increased EC_50_ value for L-glutamate, (iii) the GluN1-D732W/GluN2A and GluN1/GluN2A-S511N receptors showed regular surface expression but did not elicit any current responses in HEK293 cells, and (iv) *in silico* analysis showed that selected mutations causing structurally distinct changes in LBDs had similar effects on the surface expression of GluN1/GluN2A receptors (e.g. GluN1-D732E/GluN2A and GluN1-D732W/GluN2A receptors showed unchanged surface expression and GluN1-S688P/GluN2A and GluN1-S688C/GluN2A-receptors showed ∼59% and ∼53% reduction of surface expression).

Previous studies identified other regions in GluN subunits besides LBDs, including ATD (Qiu et al., 2009), M3 domains (Horak et al., 2008), and CTDs (Standley et al., 2000; Scott et al., 2001; Xia et al., 2001; Horak and Wenthold, 2009), containing ER retention and export signals. Therefore, we anticipate that future studies will investigate at what level other ER retention and export signals contribute to regulating the surface expression of the GluN1/GluN2 receptors with mutated LBDs. Further studies will also have to investigate the impact of the pathogenic variants in LBDs of the GluN subunits during CNS development and postnatally at the whole organism level, e.g., using knock-in mice.

## Acknowledgments

Supported by the Czech Science Foundation (24-10026S), project registration number LX22NPO5107 (MEYS CR): Financed by EU – Next Generation EU and by project registration number CZ.02.01.01/00/22_008/0004562 (Exregmed, MEYS CR). This work was also supported by the Grant Agency of Charles University (GAUK: 306221), by Healthcare Challenges of WMD II (Military Faculty of Medicine), and by the Microscopy Service Centre (Institute of Experimental Medicine CAS) supported by the MEYS CR (LM2023050; Czech-Bioimaging).

## EXTENDED FIGURE LEGENDS

**Figure 3-1.**
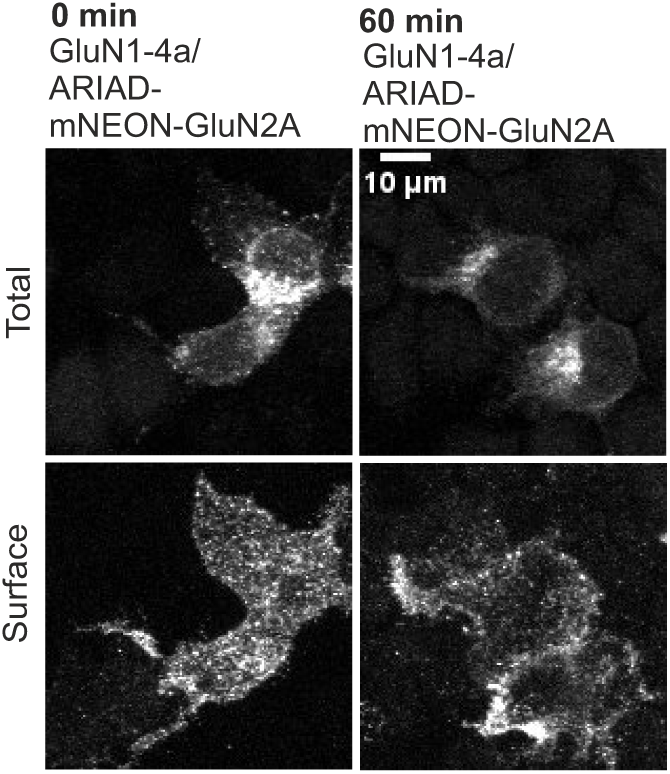
Representative images of HEK293 cells transfected with ARIAD-mNEON-GluN2A construct alone or with the GluN1-4a subunit at 0 min (without AL) and 60 min after adding AL. The total and the cell surface amount (top and bottom row, respectively) of ARIAD-mNEON-GluN2A subunits were labeled using an anti-mNEONGreen antibody 24 hours after the transfection.

**Figure 3-2.**
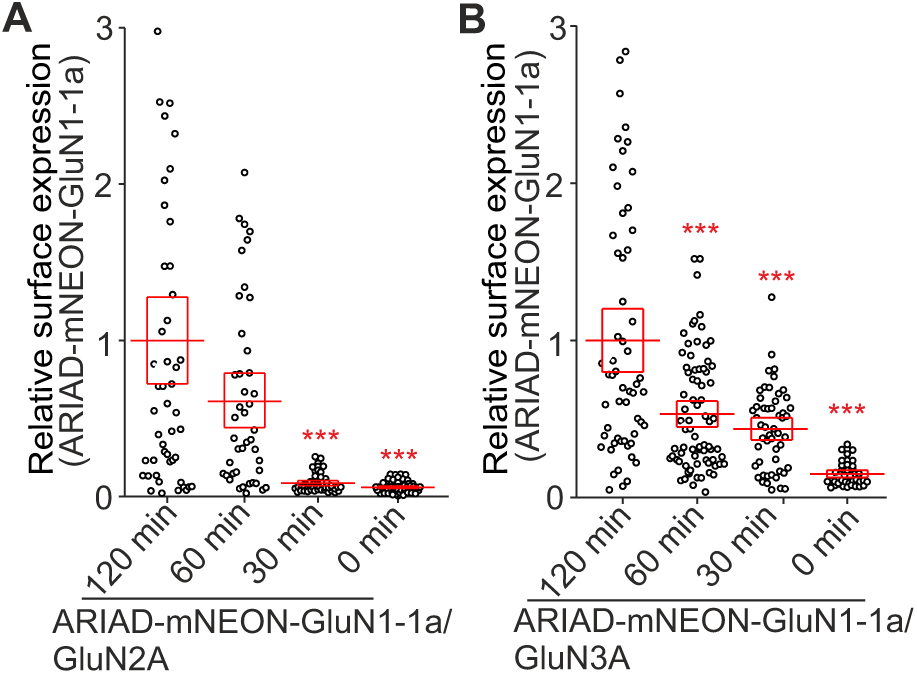
**(A,B)** Summary of relative surface expression of NMDARs containing WT ARIAD-mNEON-GluN1-1a construct together with GluN2A **(A)** or GluN3A **(B)** subunits in the presence (120, 60, 30 min) or absence (0 min) of AL. ***p < 0.001 vs 120 min of AL; one-way ANOVA. Data points correspond to individual cells (*n* ≥ 39), and the red box plot represents mean ± SEM.

**Figure 3-3.**
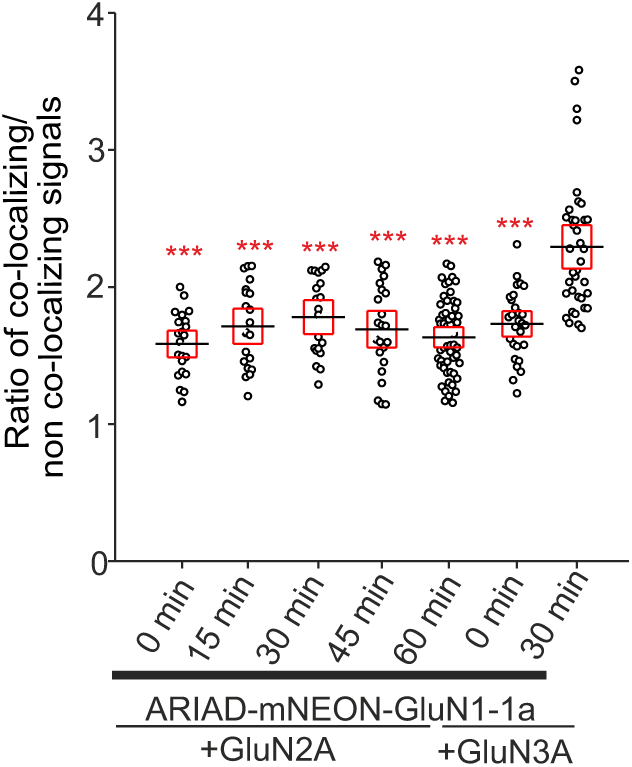
Summary of the average intensity of ARIAD-mNEON-GluN1-1a subunit signal colocalized in the presence of AL in indicated times with GM130 over the average intensity of ARIAD-mNEON-GluN1-1a subunit signal outside the GM130 signal, calculated for the indicated NMDAR combinations. \*\*\**p* < 0.001 for differences between ARIAD-mNEON-GluN1-1a/GluN3A receptor in the presence of AL vs time points for indicated NMDARs; one-way ANOVA. Data points correspond to individual cells (*n* ≥ 20), and the red box plot represents mean ± SEM.

